# Overview of the SAMPL6 p*K*_a_ Challenge: Evaluating small molecule microscopic and macroscopic p*K*_a_ predictions

**DOI:** 10.1101/2020.10.15.341792

**Authors:** Mehtap Işık, Ariën S. Rustenburg, Andrea Rizzi, M. R. Gunner, David L. Mobley, John D. Chodera

## Abstract

The prediction of acid dissociation constants (p*K*_a_) is a prerequisite for predicting many other properties of a small molecule, such as its protein-ligand binding affinity, distribution coefficient (log *D*), membrane permeability, and solubility. The prediction of each of these properties requires knowledge of the relevant protonation states and solution free energy penalties of each state. The SAMPL6 p*K*_a_ Challenge was the first time that a separate challenge was conducted for evaluating p*K*_a_ predictions as part of the Statistical Assessment of Modeling of Proteins and Ligands (SAMPL) exercises. This challenge was motivated by significant inaccuracies observed in prior physical property prediction challenges, such as the SAMPL5 log *D* Challenge, caused by protonation state and p*K*_a_ prediction issues. The goal of the p*K*_a_ challenge was to assess the performance of contemporary p*K*_a_ prediction methods for drug-like molecules. The challenge set was composed of 24 small molecules that resembled fragments of kinase inhibitors, a number of which were multiprotic. Eleven research groups contributed blind predictions for a total of 37 p*K*_a_ distinct prediction methods. In addition to blinded submissions, four widely used p*K*_a_ prediction methods were included in the analysis as reference methods. Collecting both microscopic and macroscopic p*K*_a_ predictions allowed in-depth evaluation of p*K*_a_ prediction performance. This article highlights deficiencies of typical p*K*_a_ prediction evaluation approaches when the distinction between microscopic and macroscopic p*K*_a_s is ignored; in particular, we suggest more stringent evaluation criteria for microscopic and macroscopic p*K*_a_ predictions guided by the available experimental data. Top-performing submissions for macroscopic p*K*_a_ predictions achieved RMSE of 0.7-1.0 p*K*_a_ units and included both quantum chemical and empirical approaches, where the total number of extra or missing macroscopic p*K*_a_s predicted by these submissions were fewer than 8 for 24 molecules. A large number of submissions had RMSE spanning 1-3 p*K*_a_ units. Molecules with sulfur-containing heterocycles or iodo and bromo groups were less accurately predicted on average considering all methods evaluated. For a subset of molecules, we utilized experimentally-determined microstates based on NMR to evaluate the dominant tautomer predictions for each macroscopic state. Prediction of dominant tautomers was a major source of error for microscopic p*K*_a_ predictions, especially errors in charged tautomers. The degree of inaccuracy in p*K*_a_ predictions observed in this challenge is detrimental to the protein-ligand binding affinity predictions due to errors in dominant protonation state predictions and the calculation of free energy corrections for multiple protonation states. Underestimation of ligand p*K*_a_ by 1 unit can lead to errors in binding free energy errors up to 1.2 kcal/mol. The SAMPL6 p*K*_a_ Challenge demonstrated the need for improving p*K*_a_ prediction methods for drug-like molecules, especially for challenging moieties and multiprotic molecules.

## 1 Introduction

The acid dissociation constant (*K*_a_) describes the protonation state equilibrium of a molecule given pH. More commonly, we refer to *pK*_a_ = −log_10_ ***K***_*a′*_, its negative logarithmic form. Predicting p*K*_a_ is a prerequisite for predicting many other properties of small molecules such as their protein binding affinity, distribution coefficient (log *D*), membrane permeability, and solubility. As a major aim of computer-aided drug design (CADD) is to aid in the assessment of pharmaceutical and physicochemical properties of virtual molecules prior to synthesis to guide decision-making, accurate computational p*K*_a_ predictions are required in order to accurately model numerous properties of interest to drug discovery programs.

Ionizable sites are found often in drug molecules and influence their pharmaceutical properties including target affinity, ADME/Tox, and formulation properties [1]. It has been reported that most drugs are ionized in the range of 60-90% at physiological pH [2]. Drug molecules with titratable groups can exist in many different charge and protonation states based on the pH of the environment. Given that experimental data of protonation states and p*K*_a_ are often not available, we rely on predicted p*K*_a_ values to determine which charge and protonation states the molecules populate and the relative populations of these states, so that we can assign the appropriate dominant protonation state(s) in fixed-state calculations or the appropriate solvent state weights/protonation penalty to calculations considering multiple states.

The pH of the human gut ranges between 1-8, and 74% of approved drugs can change ionization state within this physiological pH range [3]. Because of this, p*K*_a_ values of drug molecules provide essential information about their physicochemical and pharmaceutical properties. A wide distribution of acidic and basic p*K*_a_ values, ranging from 0 to 12, have been observed in approved drugs [1,3].

Drug-like molecules present difficulties for p*K*_a_ prediction compared with simple monoprotic molecules. Drug-like molecules are frequently multiprotic, have large conjugated systems, often contain heterocycles, and can tautomerize. In addition, druglike molecules with significant conformational flexibility can form intramolecular hydrogen bonding, which can significantly shift their p*K*_a_ values compared to molecules that cannot form intramolecular hydrogen bonds. This presents further challenges for modeling methods, where deficiencies in solvation models may mispredict the propensity for intramolecular hydrogen bond formation.

Accurately predicting p*K*_a_s of drug-like molecules accurately is a prerequisite for computational drug discovery and design. Small molecule p*K*_a_ predictions can influence computational protein-ligand binding affinities in multiple ways. Errors in p*K*_a_ predictions can cause modeling the wrong charge and tautomerization states which affect hydrogen bonding opportunities and charge distribution within the ligand. The dominant protonation state and relative populations of minor states in aqueous medium is dictated by the molecule’s p*K*_a_ values. The relative free energy of different protonation states in the aqueous state is a function of pH, and contributes to the overall protein-ligand affinity in the form of a free energy penalty for populating higher energy protonation states [4]. Any error in predicting the free energy of a minor aqueous protonation state of a ligand that dominates the complex binding free energy will directly add to the error in the predicted binding free energy, and selecting the incorrect dominant protonation state altogether can lead to even larger modeling errors. Similarly for log *D* predictions, an inaccurate prediction of protonation states and their relative free energies will be detrimental to the accuracy of transfer free energy predictions.

For a monoprotic weak acid (HA) or base (B)—whose dissociation equilibria are shown in Equation 1 -the acid dissociation constant is expressed as in Equation 2, or, commonly, in its negative base-10 logarithmic form as in Equation 3. The ratio of ionization states can be calculated with Henderson-Hasselbalch equations shown in Equation 4.

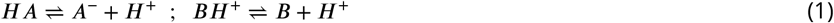

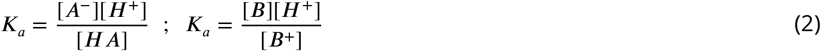

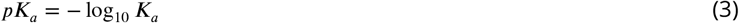

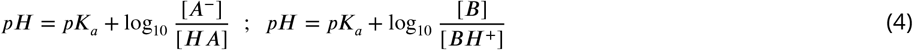

For multiprotic molecules, the definition of p*K*_a_ diverges into macroscopic p*K*_a_ and microscopic p*K*_a_ [5–7]. Macroscopic *pK_a_* describes the equilibrium dissociation constant between different charged states of the molecule. Each charge state can be composed of multiple tautomers. Macroscopic p*K*_a_ is about the deprotonation of the molecule, rather than the location of the titratable group. A microscopic p*K*_a_ describes the acid dissociation equilibrium between individual tautomeric states of different charges. (There is no p*K*_a_ defined between tautomers of the same charge as they have the same number of protons and their relative populations are independent of pH.) The microscopic p*K*_a_ determines the identity and distribution of tautomers within each charge state. Thus, each macroscopic charge state of a molecule can be composed of multiple microscopic tautomeric states. The microscopic p*K*_a_ value defined between two microstates captures the deprotonation of a single titratable group with other titratable groups held in a fixed background protonation state. In molecules with multiple titratable groups, the protonation state of one group can affect the proton dissociation propensity of another functional group, therefore the same titratable group may have different proton affinities (microscopic p*K*_a_ values) based on the protonation state of the rest of the molecule.

Different experimental methods are sensitive to changes in the total charge or the location of individual protons, so they measure different definitions of p*K*_a_s, as explained in more detail in prior work [8]. Most common p*K*_a_ measurement techniques such as potentiometric and spectrophotometric methods measure macroscopic p*K*_a_s, while NMR measurements can determine microscopic p*K*_a_s by measuring microstate populations with respect to pH. Therefore, it is important to pay attention to the source and definition of p*K*_a_ values in order to correctly interpret their meaning.

Many computational methods can predict both microscopic and macroscopic p*K*_a_s. While experimental measurements more often provide only macroscopic p*K*_a_s, microscopic p*K*_a_ predictions are more informative for determining relevant microstates (tautomers) of a molecule and their relative free energies. Predicted microstate populations can be converted to predicted macroscopic p*K*_a_s for direct comparison with experimentally obtained macroscopic p*K*_a_s. In this paper, we explore approaches to assess the performance of both macroscopic and microscopic p*K*_a_ predictions, taking advantage of available experimental data.

Microscopic p*K*_a_ predictions can be converted to macroscopic p*K*_a_ predictions either directly with Equation 5 [9],

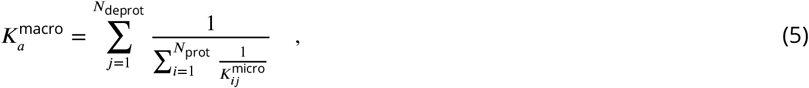

or through computing the macroscopic free energy of deprotonation between ionization states with charges *N* and *N* − 1 via Boltzmann-weighted sum of the relative free energy of microstates (*G_i_*) as in Equations 6 and 7 [10].

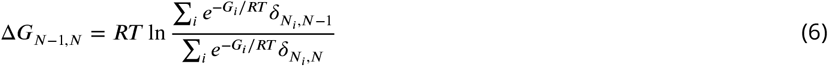

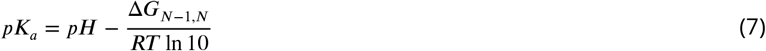

In Equation 6 Δ*G*_*N*−1, *N*_ is the effective macroscopic protonation free energy. *δ_N_i_, N−1_* is equal to unity when the microstate *i* has a total charge of *N* − 1 and zero otherwise. *RT* is the ideal gas constant times the absolute temperature.

### 1.1 Motivation for a blind p*K*_a_ challenge

SAMPL (Statistical Assessment of the Modeling of Proteins and Ligands) is a series of annual computational prediction challenges for the computational chemistry community. The goal of the SAMPL community is to evaluate the current performance of computational models and to bring the attention of the quantitative biomolecular modeling field on problems that limit the accuracy of protein-ligand binding models. SAMPL Challenges aim to enable computer-aided drug discovery to make sustained progress toward higher accuracy by focusing the community on critical challenges that isolate one accuracy-limiting problem at a time. By conducting a series of blind challenges—which often feature the computation of specific physical properties critical for protein-ligand modeling—and encouraging rapid sharing of lessons learned, SAMPL aims to accelerate progress toward quantitative accuracy in modeling.

SAMPL Challenges that focus on physical properties have assessed intermolecular binding models of various protein-ligand and host-guest systems, as well as the prediction of hydration free energies and distribution coefficients to date. These blind challenges motivate improvements in computational methods by revealing unexpected sources of error, identifying features of methods that perform well or poorly, and enabling the participants to share information after each successive challenge. Previous SAMPL Challenges have focused on the limitations of force field accuracy, finite sampling, solvation modeling defects, and tautomer/protonation state predictions on protein-ligand binding predictions.

During the SAMPL5 log *D* Challenge, the performance of models in predicting cyclohexane-water log *D* was worse than expected—accuracy suffered when protonation states and tautomers were not taken into account [11,12]. Many participants simply submitted log *P* predictions as if they were equivalent to log *D*, and many were not prepared to account for the contributions of different ionization states to the distribution coefficient in their models. Challenge results highlighted that log *P* predictions were not an accurate approximation of log *D* without capturing protonation state effects. The calculations were improved by including free energy penalty of the neutral state which relies on obtaining an accurate p*K*_a_ prediction [11]. With the goal of deconvoluting the different sources of error contributing to the large errors observed in the SAMPL5 log *D* Challenge, we organized separate p*K*_a_ and log *P* challenges in SAMPL6 [8,13,14]. For this iteration of the SAMPL challenge, we isolated the problem of predicting aqueous protonation states and associated p*K*_a_ values.

This is the first time a blind p*K*_a_ prediction challenge has been fielded as part of SAMPL. In this challenge, we aimed to assess the performance of current p*K*_a_ prediction methods for drug-like molecules, investigate potential causes of inaccurate p*K*_a_ estimates, and determine how the current level of accuracy of these models might impact the ability to make quantitative predictions of protein-ligand binding affinities.

### 1.2 Approaches to predict small molecule p*K*_a_s

There are a large variety of p*K*_a_ prediction methods developed for the prediction of aqueous p*K*_a_s of small molecules. Broadly, we can divide p*K*_a_ predictions as knowledge-based empirical methods and physical methods. Empirical methods include the following categories: Database Lookup (DL) [15], Linear Free Energy Relationship (LFER) [16–18], Quantitative Structure-Property Relationship (QSPR) [19–22], and Machine Learning (ML) approaches [23, 24]. DL methods rely on the principle that structurally similar compounds have similar p*K*_a_ values and utilize an experimental database of complete structures or fragments. The p*K*_a_ value of the most similar database entry is reported as the predicted p*K*_a_ of the query molecule. In the QSPR approach, the p*K*_a_ values are predicted as a function of various quantitative molecular descriptors, and the parameters of the function are trained on experimental datasets. A function in the form of multiple linear regression is common, although more complex forms can also be used such as the artificial neural networks in ML methods. The LFER approach is the oldest p*K*_a_ prediction strategy. They use Hammett-Taft type equations to predict p*K*_a_ based on classification of the molecule to a parent class (associated with a base p*K*_a_ value) and two parameters that describe how the base p*K*_a_ value must be modified given its substituents. Physical modeling of p*K*_a_ predictions requires Quantum Mechanics (QM) models. QM methods are often utilized together with linear empirical corrections (LEC) that are designed to rescale and unbias QM predictions for better accuracy. Classical molecular mechanics-based p*K*_a_ prediction methods are not feasible as deprotonation is a covalent bond breaking event that can only be captured by QM. Constant-pH molecular dynamics methods can calculate p*K*_a_ shifts in large biomolecular systems where there is low degree of coupling between protonation sites and linear summation of protonation energies can be assumed [25]. However, this approach can not generally be applied to small organic molecule due to the high degree of coupling between protonation sites [26–28].

## 2 Methods

### 2.1 Design and logistics of the SAMPL6 p*K*_a_ Challenge

The SAMPL6 p*K*_a_ Challenge was conducted as a blind prediction challenge and focused on predicting aqueous p*K*_a_ values of 24 small molecules not previously reported in the literature. The challenge set was composed of molecules that resemble fragments of kinase inhibitors. Heterocycles that are frequently found in FDA-approved kinase inhibitors were represented in this set. The compound selection process was described in depth in the prior publication reporting SAMPL6 p*K*_a_ Challenge experimental data collection [8]. The distribution of molecular weights, experimental p*K*_a_ values, number of rotatable bonds, and heteroatom to carbon ratio are depicted in Fig. 1. The challenge molecule set was composed of 17 small molecules with limited flexibility (less than 5 non-terminal rotatable bonds) and 7 molecules with 5-10 non-terminal rotatable bonds. The distribution of experimental p*K*_a_ values was roughly uniform between 2-12. 2D representations of all compounds are provided in Fig. 5. Drug-like molecules are often larger and more complex than the ones used in this study. We limited the size and the number of rotatable bonds of compounds to create molecule set of intermediate difficulty.

**Figure 1.**
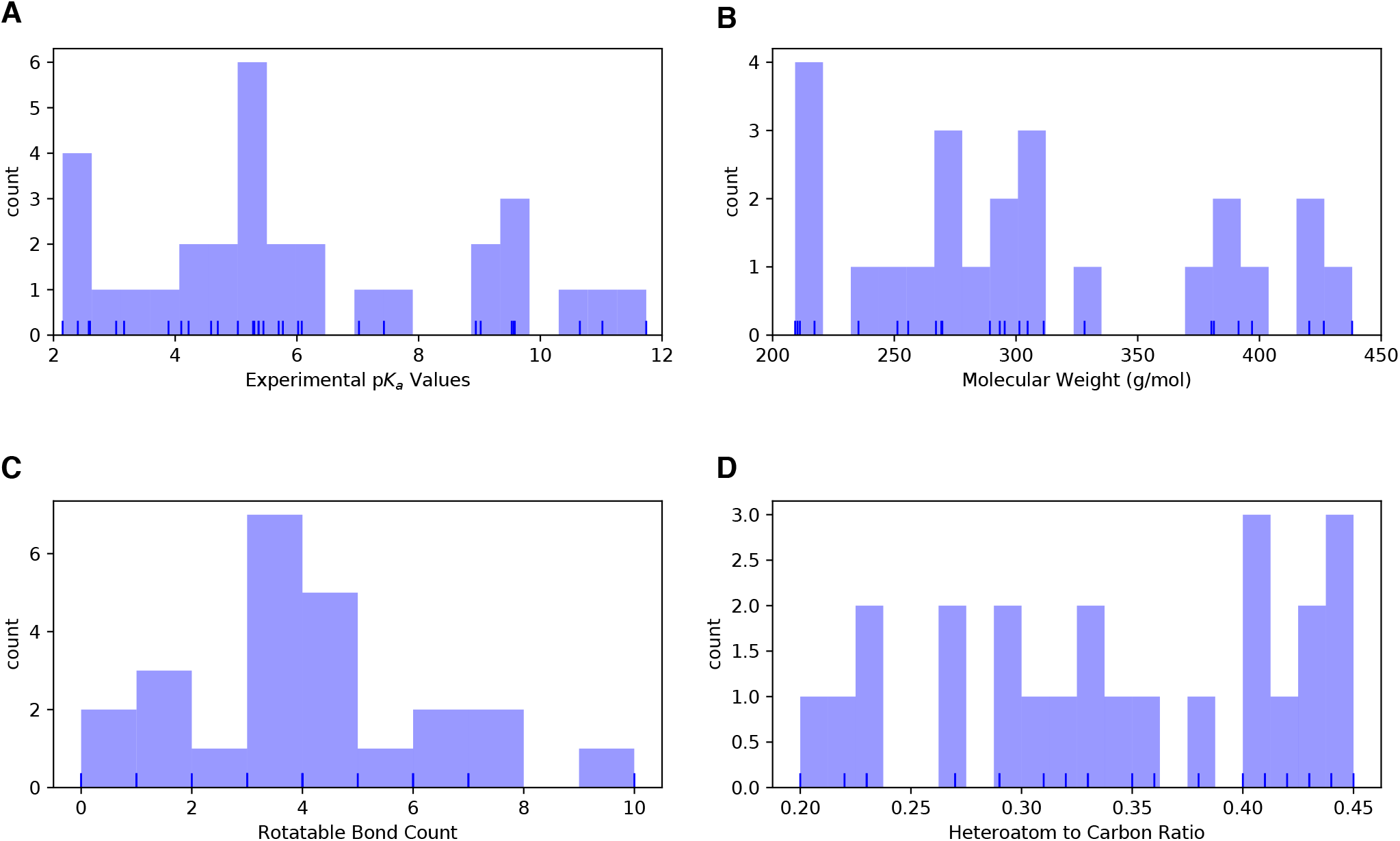
Distribution of molecular properties of the 24 compounds from the SAMPL6 p*K*_a_ Challenge. **A** Histogram of spectrophotometric p*K*_a_ measurements collected with Sirius T3 [8]. The overlaid rug plot indicates the actual values. Five compounds have multiple measured p*K*_a_s in the range of 2-12. **B** Histogram of molecular weights calculated for the neutral state of the compounds in SAMPL6 set. Molecular weights were calculated by neglecting counterions. **C** Histogram of the number of non-terminal rotatable bonds in each molecule. **D** The histogram of the ratio of heteroatom (non-carbon heavy atoms including, O, N, F, S, Cl, Br, I) count to the number of carbon atoms.

The dataset composition and experimental details—without the identity of the small molecules—were announced approximately one month before the challenge start date. Experimental macroscopic p*K*_a_ measurements were collected using a spectrophotometric method with the Sirius T3 (Sirius Analytical), at room temperature, in ionic strength-adjusted water with 0.15 M KCl [8]. The instructions for participation and the identity of the challenge molecules were released on the challenge start date (October 25, 2017). A table of molecule IDs (in the form of SM##) and their canonical isomeric SMILES was provided as input. Blind prediction submissions were accepted until January 22, 2018.

Following the conclusion of the blind challenge, the experimental data was made public on January 23, 2018. The SAMPL organizers and participants gathered at the Second Joint D3R/SAMPL Workshop at UC San Diego, La Jolla, CA on February 22–23, 2018 to share results. The workshop aimed to create an opportunity for participants to discuss the results, evaluate methodological choices by comparing the performance of different methods, and share lessons learned from the challenge. Participants reported their results and their own evaluations in a special issue of the Journal of Computer-Aided Molecular Design [29].

While designing this first p*K*_a_ prediction challenge, we did not know the optimal format to capture p*K*_a_ predictions of participants. We wanted to capture all necessary information that will aid the evaluation of p*K*_a_ predictions at the submission stage. Our strategy was to directly evaluate macroscopic p*K*_a_ predictions comparing them to experimental macroscopic p*K*_a_ values and to use collected microscopic p*K*_a_ prediction data for more in-depth diagnostics of method performance. Therefore, we asked participants to submit their predictions in three different submission types:

- **Type I:** microscopic p*K*_a_ values and related microstate pairs
- **Type II:** fractional microstate populations as a function of pH in 0.1 pH increments
- **Type III:** macroscopic p*K*_a_ values

For each submission type, a machine-readable submission file template was specified. For type I submissions, participants were asked to report the microstate ID of the protonated state, the microstate ID of deprotonated state, the microscopic p*K*_a_, and the predicted microscopic p*K*_a_ standard error of the mean (SEM). The method of microstate enumeration and why it was needed are discussed further in Section 2.2 “Enumeration of Microstates”. The SEM aims to capture the statistical uncertainty of the prediction method. Microstate IDs were preassigned identifiers for each microstate in the form of SM##_micro###. For type II submissions, the submission format included a table that started with a microstate ID column and a set of columns reporting the natural logarithm of fractional microstate population values of each predicted microstate for 0.1 pH increments between pH 2 and 12. For type III submissions participants were asked to report molecule ID, macroscopic p*K*_a_, and macroscopic p*K*_a_ SEM.

We required participants to submit predictions for all fields for each prediction, but it was not mandatory to submit predictions for all the molecules or all three submission types. Although we accepted submissions with partial sets of molecules, it would have been a better choice to require predictions for all the molecules for a better comparison of overall method performance. The submission files also included fields for naming the method, listing the software utilized, and a free text section to describe the methodology used in detail.

Participants were allowed to submit predictions for multiple methods as long as they created separate submission files. While anonymous participation was allowed, all participants opted to make their submissions public. Blind submissions were assigned a unique 5-digit alphanumeric submission ID, which will be used throughout this paper. Unique IDs were also assigned when multiple submissions exist for different submissions types of the same method such as microscopic p*K*_a_ (type I) and macroscopic p*K*_a_ (type III). These submission IDs were also reported in the evaluation papers of participants to allow cross-referencing. Submission IDs, participant-provided method names, and method categories are presented in Table 1. In many cases, multiple types of submissions (type I, II, and III) of the same method were provided by participants as challenge instructions requested. Although each prediction set was assigned a separate submission ID, we matched the submissions that originated from the same method according to the reports of the participants for cases where multiple sets of predictions came from a given method. Submission IDs for both macroscopic (type III) and microscopic (type I) p*K*_a_ predictions for each method are shown in Table 1.

**Table 1.**
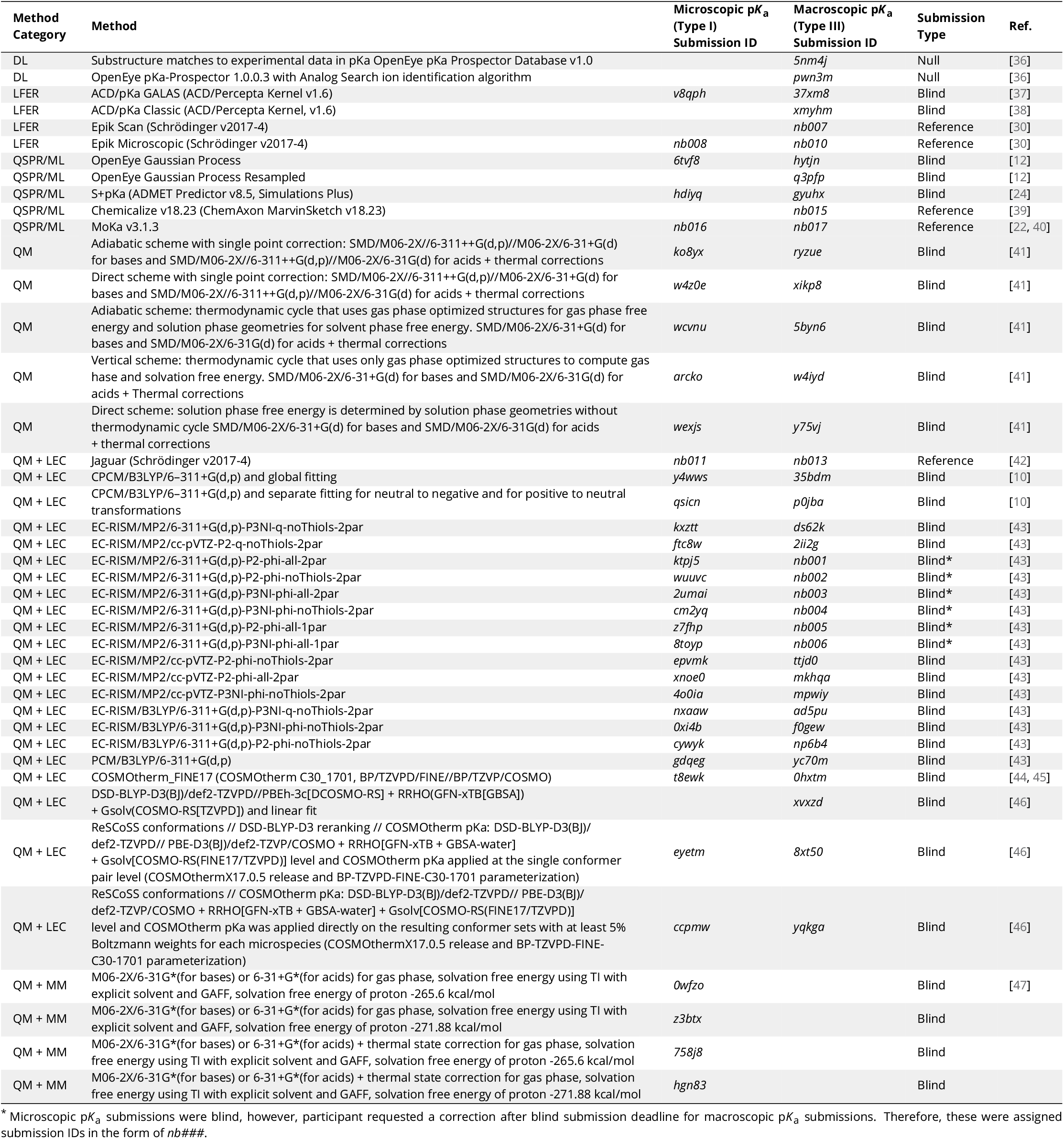
Submission IDs, names, category, and type for all the p*K*_a_ prediction sets. Reference calculations are labeled as *nb###*. The method name column lists the names provided by each participant in the submission file. The “type” column indicates if a submission was or a postdeadline reference calculation, denoted by “Blind” or “Reference” respectively. The methods in the table are grouped by method category and not ordered by performance.

### 2.2 Enumeration of microstates

To capture both the p*K*_a_ value and titrating proton position for microscopic p*K*_a_ predictions, we needed microscopic p*K*_a_ values to be reported together with a pair of microstates which describe the protonated and deprotonated states corresponding to each microscopic transition. String representations of molecules such as canonical SMILES with explicit hydrogens can be written, however, there can be inconsistencies between the interpretation of canonical SMILES written by different software and algorithms. To avoid complications while reading microstate structure files from different sources, we decided that the safest route was pre-enumerating all possible microstates of challenge compounds, assigning microstate IDs to each in the form of SM##_micro###, and requiring participants to report microscopic p*K*_a_ values along with microstate pairs specified by the provided microstates IDs.

We created initial sets of microstates with Schrödinger Epik [30] and OpenEye QUACPAC [31] and took the union of results. Microstates with Epik were generated using Schrödinger Suite v2016-4, running Epik to enumerate all tautomers within 20 p*K*_a_ units of pH 7. For enumerating microstates with OpenEye QUACPAC, we had to first enumerate formal charges and for each charge enumerate all possible tautomers using the settings of maximum tautomer count 200, level 5, with carbonyl hybridization set to False. Then we created a union of all enumerated states written as canonical isomeric SMILES generated by OpenEye OEChem [32]. Even though resonance structures correspond to different canonical isomeric SMILES, they are not different microstates, therefore it was necessary to remove resonance structures that were replicates of the same tautomer. To detect equivalent resonance structures, we converted canonical isomeric SMILES to InChI hashes with explicit and fixed hydrogen layer. Structures that describe the same tautomer but different resonance states lead to explicit hydrogen InChI hashes that are identical, allowing replicates to be removed. The Jupyter Notebook used for the enumeration of microstates is provided in Supplementary Information.

We provided microstate ID tables with canonical SMILES and 2D depictions to aid participants in matching predicted structures to microstate IDs. A canonical SMILES representation was selected over canonical isomeric SMILES, because resonance and geometric isomerism do not lead to different microstates according to our working microstate definition. The only exception was for molecule SM20, which should be consistently modeled as the E-isomer.

During the course of the SAMPL6 Challenge, participants identified new microstates that were not present in the initial list that we provided. Despite combining enumerated charge states and tautomers generated by both Epik and OpenEye QUACPAC, to our surprise, the microstate lists were still incomplete. Based on participant requests for new microstates, we iteratively had to update the list of microstates and assign new microstate IDs. Every time we received a request, we shared the updated microstate ID lists with all challenge participants. Some participants updated their p*K*_a_ prediction by including the newly added microstates in their calculations. In the future, developing a better algorithm that can enumerate all possible microstates (not just the ones with significant populations) would be very beneficial for anticipating microstates that may be predicted by p*K*_a_ prediction methods.

A microscopic p*K*_a_ definition was provided in challenge instructions for clarity as follows: Physically meaningful microscopic p*K*_a_s are defined between microstate pairs that can interconvert by single protonation/deprotonation event of only one titrable group. So, microstate pairs should have total charge (absolute) difference of 1 and only one heavy atom that differs in the number of associated hydrogens, regardless of resonance state or geometric isomerism. All geometric isomer and resonance structure pairs that have the same number of hydrogens bound to equivalent heavy atoms are grouped into the same microstate. Pairs of resonance structures and geometric isomers (cis/trans, stereo) are not considered as different microstates, as long as there is no change in the number of hydrogens bound to each heavy atom. Transitions where there are shifts in the position of protons coupled to changes in the number of protons were also not considered as microscopic p*K*_a_ values [26]. Since we wanted participants to report only microscopic p*K*_a_s that describe single deprotonation events (in contrast to transitions between microstates that are different in terms of two or more titratable protons), we have also provided a pre-enumerated list of allowed microstate pairs.

Provided microstate ID and microstate pair lists were intended to be used for reporting microstate IDs and to aid parsing of submissions. The enumerated lists of microstates were not created with the intent to guide computational predictions. This was clearly stated in the challenge instructions. However, we noticed that some participants still used the microstate lists as an input for their p*K*_a_ predictions as we received complaints from participants that due to our updates to microstate lists they needed to repeat their calculations. This would not have been an issue if participants used p*K*_a_ prediction protocols that did not rely on an external pre-enumerated list of microstates as an input. None of the participants reported this dependency in their method descriptions explicitly, so it was also not obvious how participants were using the provided states in their predictions. We could not identify which submissions used these enumerated microstate lists as input for predictions and which have followed the challenge instructions and relied only on their prediction method to generate microstates.

### 2.3 Evaluation approaches

Since the experimental data for the challenge was mainly composed of macroscopic p*K*_a_ values of both monoproticand multipro-ticcompounds, evaluation of macroscopic and microscopic p*K*_a_ predictions was not straightforward. For a subset of 8 molecules, the dominant microstate sequence could be inferred from NMR experiments. For the rest of the molecules, the only experimental information available was the macroscopic p*K*_a_ value. The experimental data—in the form of macroscopic p*K*_a_ values—did not provide any information on which group(s) are being titrated, the microscopic p*K*_a_ values, the identity of the associated macrostates (which total charge), or microstates (which tautomers). Also, experimental data did not provide any information about the charge state of protonated and deprotonated species associated with each macroscopic p*K*_a_. Typically charges of states associated with experimental p*K*_a_ values are assigned based on p*K*_a_ predictions, not experimental evidence, but we did not utilize such computational charge assignment. For a fair performance comparison between methods, we avoided relying on any particular p*K*_a_ prediction to assist the interpretation of the experimental reference data. This choice complicated the p*K*_a_ prediction analysis, especially regarding how to pair experimental and predicted p*K*_a_ values for error analysis. We adopted various evaluation strategies guided by the experimental data. To compare macroscopic p*K*_a_ predictions to experimental values, we had to utilize numerical matching algorithms before we could calculate performance statistics. For the subset of molecules with experimental data about microstates, we used microstate-based matching. These matching methods are described in more detail in the next section.

Three types of submissions were collected during the SAMPL6 p*K*_a_ Challenge. We have only utilized the type I (microscopic p*K*_a_ value and microstate IDs) and the type III (macroscopic p*K*_a_ value) predictions in this article. Type I submissions contained the same prediction information as the type II submissions which reported the fractional population of microstates with respect to pH. We collected type II submissions in order to capture relative populations of microstates, not realizing they were redundant. The microscopic p*K*_a_ predictions collected in type I submissions capture all the information necessary to calculate type II submissions. Therefore, we did not use type II submissions for challenge evaluation. In theory, type III (macroscopic p*K*_a_) predictions can also be calculated from type I submissions, but collecting type III submissions allowed the participation of p*K*_a_ prediction methods that directly predict macroscopic p*K*_a_ values without considering microspeciation and methods that apply special empirical corrections for macroscopic p*K*_a_ predictions.

#### 2.3.1 Matching algorithms for pairing predicted and experimental p*K*_a_ values

Macroscopic p*K*_a_ predictions can be calculated from microscopic p*K*_a_ values for direct comparison to experimental macroscopic p*K*_a_ values. One major question must be answered to allow this comparison: How should we match predicted macroscopic p*K*_a_ values to experimental macroscopic p*K*_a_ values when there could multiple p*K*_a_ values reported for a given molecule? For example, experiments on SM18 showed three macroscopic p*K*_a_s, but prediction of *xvxzd* method reported two macroscopic p*K*_a_ values. There were also examples of the opposite situation with more predicted p*K*_a_ values than experimentally determined macroscopic p*K*_a_s: One experimental p*K*_a_ was measured for SM02, but two macroscopic p*K*_a_ values were predicted by *xvxzd* method. The experimental and predicted values must be paired before any prediction error can be calculated, even though there was not any experimental information regarding underlying tautomer and charge states.

Knowing the charges of macrostates would have guided the pairing between experimental and predicted macroscopic p*K*_a_ values, however, not all experimental p*K*_a_ measurements can determine determine the charge of protonation states. The potentiometric p*K*_a_ measurements just captures the relative charge change between macrostates, but not the absolute value of the charge. Thus, our experimental data did not provide any information that would indicate the titration site, the overall charge, or the tautomer composition of macrostate pairs that are associated with each measured macroscopic p*K*_a_ that can guide the matching between predicted and experimental p*K*_a_ values.

For evaluating macroscopic p*K*_a_ predictions taking the experimental data as reference, Fraczkiewicz [23] delineated recommendations for fair comparative analysis of computational p*K*_a_ predictions. They recommended that, in the absence of any experimental information that would aid in matching, experimental and computational p*K*_a_ values should be matched preserving the order of p*K*_a_ values and minimizing the sum of absolute errors.

We picked the Hungarian matching algorithm [33, 34] to match experimental and predicted macroscopic p*K*_a_ values with a squared error cost function as suggested by Kiril Lanevskij via personal communication. The algorithm is available in the SciPy package (*scipy.optimize.linear_sum_assignment*) [35]. This matching algorithm provides optimum global assignment that minimizes the linear sum of squared errors of all pairwise matches. We selected the squared error cost function instead of the absolute error cost function to avoid misordered matches, For instance, for a molecule with experimental p*K*_a_ values of 4 and 6, and predicted p*K*_a_ values of 7 and 8, Hungarian matching with absolute error cost function would match 6 to 7 and 4 to 9. Hungarian matching with squared error cost would match 4 to 7 and 6 to 9, preserving the increasing p*K*_a_ value order between experimental and predicted values. A weakness of this approach would be failing to match the experimental value of 6 to predicted value of 7 if that was the correct match based on underlying macrostates. But the underlying pair of states were unknown to us both because the experimental data did not determine which charge states the transitions were happening between and also because we did not collect the pair of macrostates associated with each p*K*_a_ predictions in submissions. Requiring this information for macroscopic p*K*_a_ predictions in future SAMPL challenges would allow for better comparison between predictions, even if experimental assignment of charges is not possible. There is no perfect solution to the numerical p*K*_a_ assignment problem, but we tried to determine the fairest way to penalize predictions based on their numerical deviation from the experimental values.

For the analysis of microscopic p*K*_a_ predictions we adopted a different matching approach. For the eight molecules for which we had the requisite data for this analysis, we utilized the dominant microstate sequence inferred from NMR experiments to match computational predictions and experimental p*K*_a_ values. We will refer to this assignment method as microstate matching, where the experimental p*K*_a_ value is matched to the computational microscopic p*K*_a_ value which was reported for the dominant microstate pair observed for each transition. We have compared the results of Hungarian matching and microstate matching.

Inevitably, the choice of matching algorithms to assign experimental and predicted values has an impact on the computed performance statistics. We believe the Hungarian algorithm for numerical matching of unassigned p*K*_a_ values and microstatebased matching when experimental microstates are known were the best choices, providing the most unbiased matching without introducing assumptions outside of the experimental data.

#### 2.3.2 Statistical metrics for submission performance

A variety of accuracy and correlation statistics were considered for analyzing and comparing the performance of prediction methods submitted to the SAMPL6 p*K*_a_ Challenge. Calculated performance statistics of predictions were provided to participants before the workshop. Details of the analysis and scripts are maintained on the SAMPL6 GitHub Repository (described in Section 5).

##### Error metrics

There are six error metrics reported for the numerical error of the p*K*_a_ values: the root-mean-squared error (RMSE), mean absolute error (MAE), mean error (ME), coefficient of determination (R^2^), linear regression slope (m), and Kendall’s Rank Correlation Coefficient *(**τ***). Uncertainty in each performance statistic was calculated as 95% confidence intervals estimated by non-parametric bootstrapping (sampling with replacement) over predictions with 10 000 bootstrap samples. Calculated errors statistics of all methods can be found in Table S2 for macroscopic p*K*_a_ predictions and Tables S4 and S4 for microscopic p*K*_a_ predictions.

##### Assessing macrostate predictions

In addition to assessing the numerical error in predicted p*K*_a_ values, we also evaluated predictions in terms of their ability to capture the correct macrostates (ionization states) and microstates (tautomers of each ionization state) to the extent possible from the available experimental data. For macroscopic p*K*_a_s, the spectrophotometric experiments do not directly report on the identity of the ionization states. However, the number of ionization states indicates the number of macroscopic p*K*_a_s that exists between the experimental range of 2.0-12.0. For instance, SM14 has two experimental p*K*_a_s and therefore three different charge states observed between pH 2.0 and 12.0. If a prediction reported 4 macroscopic p*K*_a_s, it is clear that this method predicted an extra ionization state. With this perspective, we reported the number of unmatched experimental p*K*_a_s (the number of missing p*K*_a_ predictions, i.e., missing ionization states) and the number of unmatched predicted p*K*_a_s (the number of extra p*K*_a_ predictions, i.e., extra ionization states) after Hungarian matching. The latter count was restricted to only predictions with p*K*_a_ values between 2 and 12 because that was the range of the experimental method. Errors in extra or missing p*K*_a_ prediction errors highlight failure to predict the correct number of ionization states within a pH range.

##### Assessing microstate predictions

For the evaluation of microscopic p*K*_a_ predictions, taking advantage of the available dominant microstate sequence data for a subset of 8 compounds, we calculated the dominant microstate prediction accuracy which is the ratio of correct dominant tautomer predictions for each charge state divided by the total number of dominant tautomer predictions. Dominant microstateprediction accuracy was calculated over all experimentally detected ionization states of each molecule which were part of this analysis. In order to extract the sequence of dominant microstates from the microscopic p*K*_a_ predictions sets, we calculated the relative free energy of microstates selecting a neutral tautomer and pH 0 as reference following Equation 8. Calculation of relative microstate free energies was explained in more detail in a previous publication [26].

The relative free energy of a state with respect to reference state B at pH 0.0 (arbitrary pH value selected as reference) can be calculated as follows:

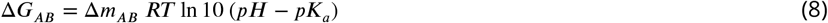

Δ*m_AB_* is equal to the number protons in state A minus that in state B. R and T indicate the molar gas constant and temperature, respectively. By calculating relative free energies of all predicted microstates with respect to the same reference state and pH, we were able to determine the sequence of predicted dominant microstates. The dominant tautomer of each charge state was determined as the microstate with the lowest free energy in the subset of predicted microstates of each ionization state. This approach is feasible because the relative free energy of tautomers of the same ionization state is independent of pH and therefore the choice of reference pH is arbitrary.

##### Identifying consistently top-performing methods

We created a shortlist of top-performing methods for macroscopic and microscopic p*K*_a_ predictions. The top macroscopic p*K*_a_ predictions were selected if they ranked in the top 10 consistently according to two error metrics (RMSE, MAE) and two correlation metrics (R-Squared, and Kendall’s Tau), while also having fewer than eight missing or extra macroscopic p*K*_a_s for the entire molecule set (eight macrostate errors correspond to macrostate prediction mistake in roughly one third of the 24 compounds). These methods are presented in Table 2. A separate list of top-performing methods was constructed for microscopic p*K*_a_ with the following criteria: ranking in the top 10 methods when ranked by accuracy statistics (RMSE and MAE) and perfect dominant microstate prediction accuracy. These methods are presented in Table 3.

**Table 2.** Four consistently well-performing prediction methods for macroscopic p*K*_a_ prediction based on consistent ranking within the Top 10 according to various statistical metrics. Submissions were ranked according to RMSE, MAE, R^2^, and *τ*. Consistently wellperforming methods were selected as the ones that rank in the Top 10 in each of these statistical metrics. These methods also have less than 2 unmatched experimental p*K*_a_s and less than 7 unmatched predicted p*K*_a_s according to Hungarian matching. Performance statistics are provided as mean and 95% confidence intervals.

| Submission ID | Method Name | RMSE | MAE | $R^2$ | Kendall's Tau ($\tau$) | Unmatched Exp. $pK_a$ Count | Unmatched Pred. $pK_a$ Count [2,12] |
| --- | --- | --- | --- | --- | --- | --- | --- |
| *xvxzd* | Full quantum chemical calculation of free energies and fit to experimental $pK_a$ | 0.68 [0.54, 0.81] | 0.58 [0.45, 0.71] | 0.94 [0.88, 0.97] | 0.82 [0.68, 0.92] | 2 | 4 |
| *gyuhx* | S+$pK_a$ | 0.73 [0.55, 0.91] | 0.59 [0.44, 0.74] | 0.93 [0.88, 0.96] | 0.88 [0.8, 0.94] | 0 | 7 |
| *xmyhm* | ACD/$pK_a$ Classic | 0.79 [0.52, 1.03] | 0.56 [0.38, 0.77] | 0.92 [0.85, 0.97] | 0.81 [0.68, 0.9] | 0 | 3 |
| *8xt50* | ReSCoSS conformations // DSD-BLYP-D3 reranking // COSMOtherm $pK_a$ | 1.07 [0.78, 1.36] | 0.81 [0.58, 1.07] | 0.91 [0.84, 0.95] | 0.80 [0.68, 0.89] | 0 | 0 |

**Table 3.** Top-performing methods for microscopic p*K*_a_ predictions based on consistent ranking within the Top 10 according to various statistical metrics calculated for 8 molecule dataset. Performance statistics are provided as mean and 95% confidence intervals. Submissions that rank in the Top 10 according to RMSE and MAE and have perfect dominant microstate prediction accuracy were selected as consistently well-performing methods. Correlation-based statistics (R^2^, and Kendall’s Tau), although reported in the table, were excluded from the statistics used for determining top-performing methods. This was because correlation-based statistics were not very discriminating due to the narrow dynamic range and the small number of data points in the 8 molecule dataset with NMR-determined dominant microstates.

| Submission ID | Method Name | Dominant Microstate Accuracy | RMSE | MAE | $R^2$ | Kendall's Tau | Unmatched Exp. $pK_a$ Count | Unmatched Pred. $pK_a$ Count [2,12] |
| --- | --- | --- | --- | --- | --- | --- | --- | --- |
| nb016 | MoKa | 1.0 [1.0, 1.0] | 0.52 [0.25, 0.71] | 0.43 [0.23, 0.65] | 0.92 [0.05, 0.99] | 0.62 [-0.14, 1.00] | 0 | 3 |
| hdiyq | S+pKa | 1.0 [1.0, 1.0] | 0.68 [0.49, 0.83] | 0.60 [0.39, 0.80] | 0.86 [0.47, 0.98] | 0.78 [0.40, 1.00] | 0 | 16 |
| nb011 | Jaguar | 1.0 [1.0, 1.0] | 0.72 [0.35, 1.07] | 0.54 [0.28, 0.86] | 0.86 [0.18, 0.98] | 0.64 [0.26, 0.95] | 0 | 36 |
| 6tvf8 | OE Gaussian Process | 1.0 [1.0, 1.0] | 0.76 [0.55, 0.95] | 0.68 [0.46, 0.90] | 0.92 [0.78, 0.99] | 0.87 [0.6, 1.00] | 0 | 55 |
| 0xi4b | EC-RISM/B3LYP/6-311+G(d,p)<br>-P3NI-phi-noThiols-2par | 1.0 [1.0, 1.0] | 1.15 [0.75, 1.50] | 0.98 [0.63, 1.36] | 0.77 [0.02, 0.98] | 0.51 [-0.14, 1.00] | 0 | 33 |
| cywyk | EC-RISM/B3LYP/6-311+G(d,p)<br>-P2-phi-noThiols-2par | 1.0 [1.0, 1.0] | 1.17 [0.88, 1.41] | 1.06 [0.74, 1.35] | 0.73 [0.02, 0.98] | 0.56 [-0.08, 1.00] | 0 | 36 |

##### Determining challenging molecules

In addition to comparing the performance of methods, we also wanted to compare p*K*_a_ prediction performance for each molecule to determine which molecules were the most challenging for p*K*_a_ predictions considering all the methods in the challenge. For this purpose, we plotted prediction error distributions of each molecule calculated over all prediction methods. We also calculated MAE for each molecule over all prediction sets as well as for predictions from each method category separately.

### 2.4 Reference calculations

Including a null model is helpful in comparative performance analysis of predictive methods to establish what the performance statistics look like for a baseline method for the specific dataset. Null models or null predictions employ a simple prediction model which is not expected to be particularly successful, but it provides a simple point of comparison for more sophisticated methods. The expectation or goal is for more sophisticated or costly prediction methods to outperform the predictions from a null model, otherwise the simpler null model would be preferable. In SAMPL6 p*K*_a_ Challenge there were two blind submissions using database lookup methods that were submitted to serve as null predictions. These methods, with submission IDs *5nm4j* and *5nm4j* both used OpenEye pKa-Prospector database to find the most similar molecule to query molecule and simply reported its p*K*_a_ as the predicted value. Database lookup methods with a rich experimental database do present a challenging null model to beat, however, due to the accuracy level needed from p*K*_a_ predictions for computer-aided drug design we believe such methods provide an appropriate performance baseline that physical and empirical p*K*_a_ prediction methods should strive to outperform.

We also included additional reference calculations in the comparative analysis to provide more perspective. Some widely used methods by academia and industry were missing from the blind challenge submission. Therefore, we included those methods as reference calculations: Schrödinger/Epik(*nb007, nb008, nb010*), Schrödinger/Jaguar(*nb011, nb013*), Chemaxon/Chemicalize (*nb015*), and Molecular Discovery/MoKa (*nb016, nb017*). Epik and Jaguar p*K*_a_ predictions were collected by Bas Rustenburg, Chemicalize predictions by Mehtap Isik, and MoKa predictions by Thomas Fox. All were done after the challenge deadline avoiding any alterations to their respective standard procedures and any guidance from experimental data. Experimental data was publicly available before these calculations were complete, therefore reference calculations were not formally considered as blind submissions.

All figures and statistics tables in this manuscript include reference calculations. As the reference calculations were not formal submissions, these were omitted from formal ranking in the challenge, but we present plots in this article which show them for easy comparison. These are labeled with submission IDs of the form *nb###* to clearly indicate non-blind reference calculations.

## 3 Results and Discussion

Participation in the SAMPL6 p*K*_a_ Challenge was high with 11 research groups contributing p*K*_a_ prediction sets for 37 methods. A large variety of p*K*_a_ prediction methods were represented in the SAMPL6 Challenge. We categorized these submissions into four method classes: database lookup (DL), linear free energy relationship (LFER), quantitative structure-property relationship or machine learning (QSPR/ML), and quantum mechanics (QM). Quantum mechanics models were subcategorized into QM methods with and without linear empirical correction (LEC), and combined quantum mechanics and molecular mechanics (QM + MM). Table 1 presents method names, submission IDs, method categories, and also references for each approach. Integral equation-based approaches (e.g.EC-RISM) were also evaluated under the Physical (QM) category. There were 2 DL, 4 LFER, and 5 QSPR/ML methods represented in the challenge, including the reference calculations. The majority of QM calculations include linear empirical corrections (22 methods in QM + LEC category), and only 5 QM methods were submitted without any empirical corrections. There were 4 methods that used a mixed physical modeling approach of QM + MM.

The following sections present a detailed performance evaluation of blind submissions and reference prediction methods for macroscopic and microscopic p*K*_a_ predictions. Performance statistics of all the methods can be found in Tables S2 and S4. Methods are referred to by their submission ID’s which are provided in Table 1.

### 3.1 Analysis of macroscopic p*K*_a_ predictions

The performance of macroscopic p*K*_a_ predictions was analyzed by comparison to experimental p*K*_a_ values collected by the spectrophotometric method via numerical matching following the Hungarian method. Overall p*K*_a_ prediction performance was worse than we hoped. Fig. 2 shows RMSE calculated for each prediction method represented by their submission IDs. Other performance statistics are depicted in Fig. 3. In both figures, method categories are indicated by the color of the error bars. The statistics depicted in these figures can be found in Table S2. Prediction error ranged between 0.7 to 3.2 p*K*_a_ units in terms of RMSE, while an RMSE between 2-3 log units was observed for the majority of methods (20 out of 38 methods). Only five methods achieved RMSE less than 1 p*K*_a_ unit. One is QM method with COSMO-RS approach for solvation and linear empirical correction (*xvxzd* (DSD-BLYP-D3(BJ)/def2-TZVPD//PBEh-3c[DCOSMO-RS] + RRHO(GFN-xTB[GBSA]) + Gsolv(COSMO-RS[TZVPD]) and linear fit)), and the remaining four are empirical prediction methods of LFER (*xmyhm* (ACD/pKa Classic), *nb007* (Schrödinger/Epik Scan)) and QSPR/ML categories (*gyuhx* (Simulations Plus), *nb017* (MoKa)). These five methods with RMSE less than 1 p*K*_a_ unit are also the methods that have the lowest MAE. *xmyhm* and *xvxzd* were the only two methods for which the upper 95% confidence interval of RMSE was lower than 1 p*K*_a_ unit.

**Figure 2.**
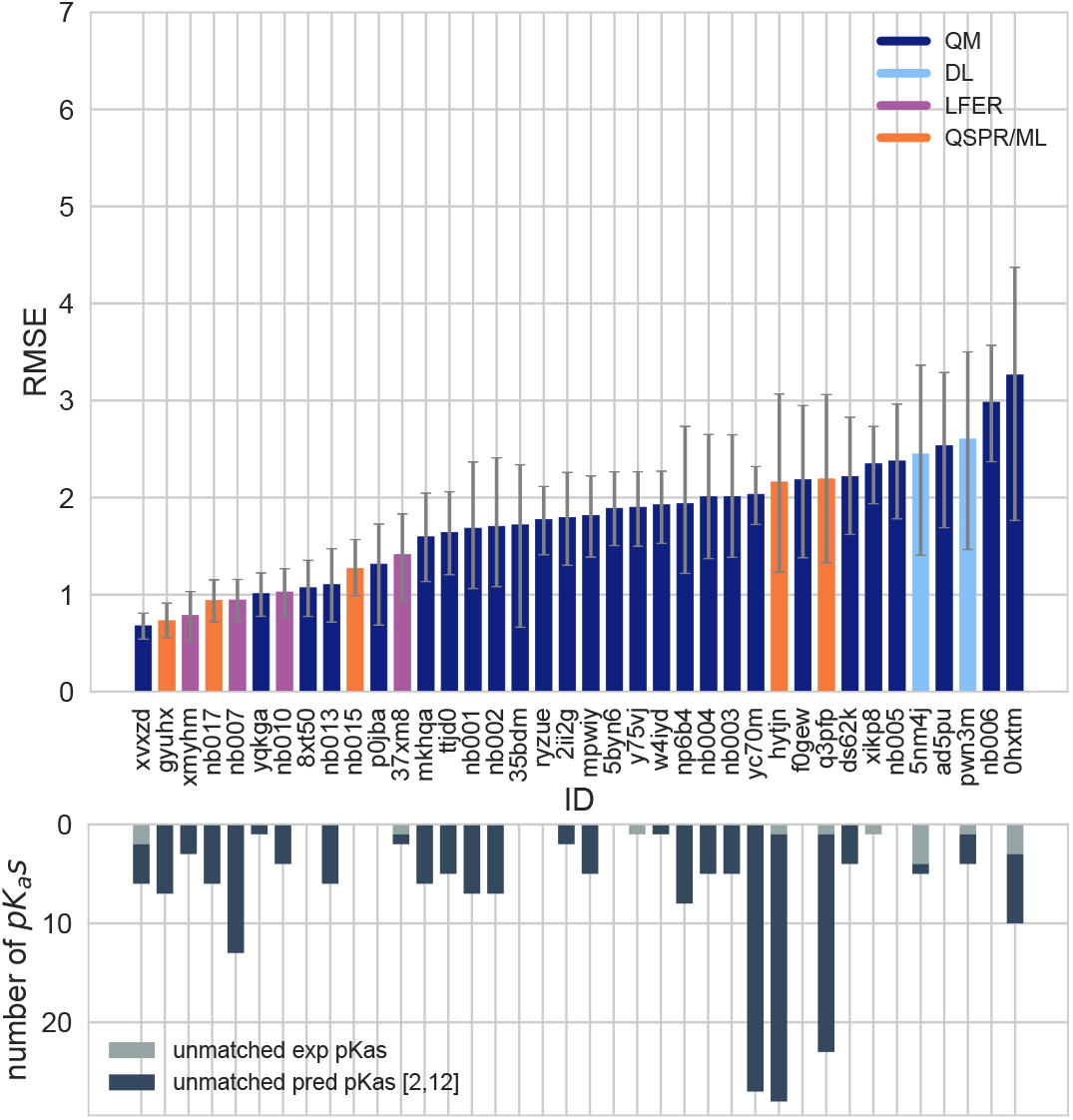
RMSE and unmatched p*K*_a_ counts vs. submission ID plots for macroscopic p*K*_a_ predictions based on Hungarian matching. Methods are indicated by submission IDs. RMSE is shown with error bars denoting 95% confidence intervals obtained by bootstrapping over challenge molecules. Submissions are colored by their method categories. Light blue colored database lookup methods are utilized as the null prediction method. QM methods category (navy) includes pure QM, QM+LEC, and QM+MM approaches. Lower bar plots show the number of unmatched experimental p*K*_a_ values (light grey, missing predictions) and the number of unmatched p*K*_a_ predictions (dark grey, extra predictions) for each method between pH 2 and 12. Submission IDs are summarized in Table 1. Submission IDs of the form *nb###* refer to non-blinded reference methods computed after the blind challenge submission deadline. All others refer to blind, prospective predictions.

**Figure 3.**
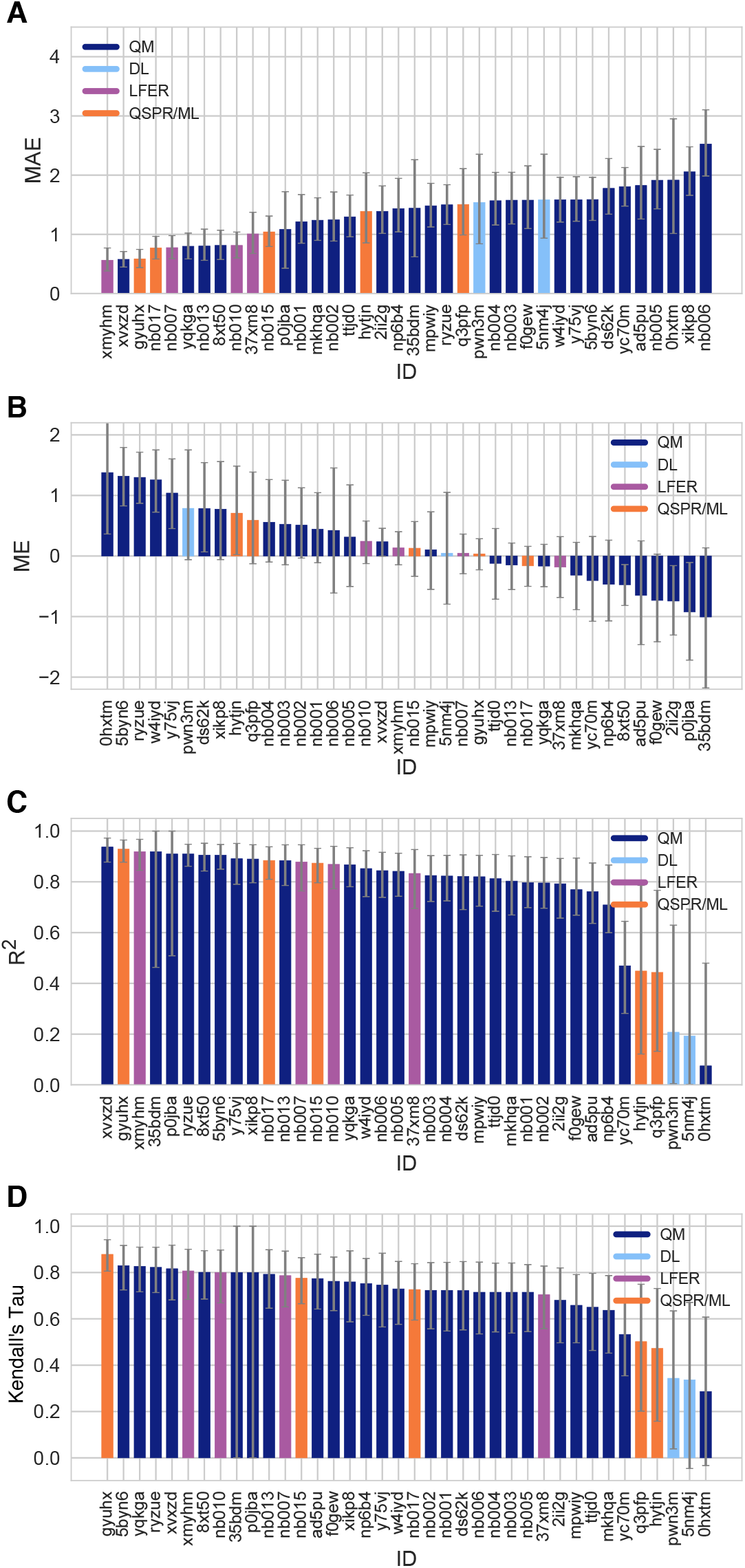
Additional performance statistics for macroscopic p*K*_a_ predictions based on Hungarian matching. Methods are indicated by submission IDs. Mean absolute error (MAE), mean error (ME), Pearson’s R^2^, and Kendall’s Rank Correlation Coefficient Tau (*τ*) are shown, with error bars denoting 95% confidence intervals were obtained by bootstrapping over challenge molecules. Refer to Table 1 for the submission IDs and method names. Submissions are colored by their method categories. Light blue colored database lookup methods are utilized as the null prediction method.

In terms of correlation statistics, many methods have good performance, although the ranking of methods changes according to R^2^ and Kendall’s Tau. Therefore, many methods are indistinguishable from one another, considering the uncertainty of the correlation statistics. 32 out of 38 methods have R and Kendall’s Tau higher than 0.7 and 0.6, respectively. 8 methods have R^2^ higher than 0.9 and 6 methods have Kendall’s Tau higher than 0.8. The overlap of these two sets are the following: *gyuhx* (Simulations Plus), *xvxzd* (DSD-BLYP-D3(BJ)/def2-TZVPD//PBEh-3c[DCOSMO-RS] + RRHO(GFN-xTB[GBSA]) + Gsolv(COSMO-RS[TZVPD]) and linear fit), *xmyhm* (ACD/pKa Classic), *ryzue* (Adiabatic scheme with single point correction: MD/M06-2X//6-311++G(d,p)//M06-2X/6-31+G(d) for bases and SMD/M06-2X//6-311++G(d,p)//M06-2X/6-31G(d) for acids + thermal corrections), and *5byn6* (Adiabatic scheme: thermodynamic cycle that uses gas phase optimized structures for gas phase free energy and solution phase geometries for solvent phase free energy. SMD/M06-2X/6-31+G(d)for bases and SMD/M06-2X/6-31G(d) for acids + thermal corrections). It is worth noting that *ryzue* and *5byn6* are QM predictions without any empirical correction. Their high correlation and rank correlation coefficient scores signal that with an empirical correction their accuracy based performance could improve. Indeed, the participants have shown that this is the case in their own challenge analysis paper and achieved RMSE of 0.73 p*K*_a_ units after the challenge [41].

Null prediction methods based on database lookup (*5nm4j* and *pwn3m*) had similar performance, with an RMSE of roughly 2.5 p*K*_a_ units, an MAE of 1.5 p*K*_a_ units, R^2^ of 0.2, and Kendall’s Tau of 0.3. Many methods were observed to have a prediction performance advantage over the null predictions shown in light blue in Fig. 2 and Fig. 3 considering all the performance metrics as a whole. In terms of correlation statistics, the null methods are the worst performers, except for *0hxtm*. From the perspective of accuracy-based statistics (RMSE and MAE), only the top 10 methods were observed to have significantly lower errors than the null methods considering the uncertainty of error metrics expressed as 95% confidence intervals.

The distribution of macroscopic p*K*_a_ prediction signed errors observed in each submission was plotted in Fig. 7A as ridge plots using the Hungarian matching scheme. *2ii2g, f0gew, np64b, p0jba*, and *yc70m* tended to overestimate, while *5byn6, ryzue*, and *w4iyd* tended to underestimate macroscopic p*K*_a_ values.

Four submissions in the QM+LEC category used the COSMO-RS implicit solvation model. While three of these achieved the lowest RMSE among QM-based methods (*xvxzd*, *yqkga*, and *8xt50*) [46], one of them showed the highest RMSE (*0hxtm* (COSMOtherm_FINE17)) among all SAMPL6 Challenge macroscopic p*K*_a_ predictions. All four methods used COSMO-RS/FINE17 to compute solvation free energies. The major difference between the three low-RMSE methods and *0hxtm* seems to be the protocol for determining relevant conformations for each microstate. *xvxzd,yqkga*, and *8xt50* used a semi-empirical tight binding (GFN-xTB) method and GBSA continuum solvation model for geometry optimization, followed by high level single-point energy calculations with a solvation free energy correction (COSMO-RS(FINE17/TZVPD)) and rigid rotor harmonic oscillator (RRHO[GFN-xTB(GBSA]) correction. *yqkga*, and *8xt50* selected conformations for each microstate with the Relevant Solution Conformer Sampling and Selection (ReSCoSS) workflow [46]. The conformations were clustered according to shape, and the lowest energy conformations from each cluster (according to BP86/TZVP/COSMO single point energies in any of the 10 different COSMO-RS solvents) were considered as relevant conformers. The*yqkga* method further filtered out conformers that have less than 5% Boltzmann weights at the DSD-BLYP-D3/def2-TZVPD + RRHO(GFNxTB) + COSMO-RS(fine) level. The *xvxzd* method used an MF-MD-GC//GFN-xTB workflow and energy thresholds of 6 kcal/mol and 10 kcal/mol, for conformer and microstate selection. On the other hand, the conformational ensemble captured for each microstate seems to be more limited for the *0hxtm* method, judging by the method description provided in the submission file (this participant did not publish an analysis of the results that they obtained for SAMPL6). The *0hxtm* method reported that relevant conformations were computed with the COSMOconf 4.2 workflow which produced multiple relevant conformers for only the neutral states of SM18 and SM22. In contrast to *xvxzd,yqkga*, and *8xt50*, the *0hxtm* method also did not include a RRHO correction. Participants who submitted the three low-RMSE methods report that capturing the chemical ensemble for each molecule including conformers and tautomers and high-level QM calculations led to more successful macroscopic p*K*_a_ prediction results and RRHO correction provided a minor improvement [46]. Comparing these results to other QM approaches in the SAMPL Challenge also points to the advantage of the COSMO-RS solvation approach compared to other implicit solvent models.

In addition to the statistics related to the p*K*_a_ value, we also analyzed missing or extra p*K*_a_ predictions. Analysis of the p*K*_a_ values with accuracy- and correlation-based error metrics was only possible after the matching of predicted macroscopic p*K*_a_ values to experimental p*K*_a_ values through Hungarian matching, although this approach masks p*K*_a_ prediction issues in the form of extra or missing macroscopic p*K*_a_ predictions. To capture this class of prediction errors, we reported the number of unmatched experimental p*K*_a_s (missing p*K*_a_ predictions) and the number of unmatched predicted p*K*_a_s (extra p*K*_a_ predictions) after Hungarian matching for each method. Both missing and extra p*K*_a_ prediction counts were only considered for the pH range of 2-12, which corresponds to the limits of the experimental assay. The lower subplot of Fig. 2 shows the total count of unmatched experimental or predicted p*K*_a_ values for all the molecules in each prediction set. The order of submission IDs in the x-axis follows the RMSD based ranking so that the performance of each method from both p*K*_a_ value accuracy and the number of p*K*_a_s can be viewed together. The omission or inclusion of extra macroscopic p*K*_a_ predictions is a critical error because inaccuracy in predicting the correct number of macroscopic transitions shows that methods are failing to predict the correct set of charge states, i.e., failing to predict the correct number of ionization states that can be observed between the specified pH range.

In the analysis of these challenge results, extra macroscopic p*K*_a_ predictions were found to be more common than missing p*K*_a_ predictions. In p*K*_a_ prediction evaluations, the accuracy of predicted ionization states within a pH range is usually neglected. When predictions are only evaluated for the accuracy of the p*K*_a_ value with numerical matching algorithms, a larger number of predicted p*K*_a_s lead to greater underestimation of prediction errors. Therefore, it is not surprising that methods are biased to predict extra p*K*_a_ values. The SAMPL6 p*K*_a_ Challenge experimental data consists of 31 macroscopic p*K*_a_s in total, measured for 24 molecules (6 molecules in the set have multiple p*K*_a_s). Within the 10 methods with the lowest RMSE, only the *xvxzd* method predicts too few p*K*_a_ values (2 unmatched out of 31 experimental p*K*_a_s). All other methods that rank in the top 10 by RMSE have extra predicted p*K*_a_s ranging from 1 to 13. Two prediction sets without any extra p*K*_a_ predictions and low RMSE are *8xt50* (ReSCoSS conformations // DSD-BLYP-D3 reranking // COSMOtherm pKa) and *nb015* (ChemAxon/Chemicalize).

#### 3.1.1 Consistently well-performing methods for macroscopic p*K*_a_ prediction

Methods ranked differently when ordered by different error metrics, although there were a couple of methods that consistently ranked in the top fraction. By using combinatorial criteria that take multiple statistical metrics and unmatched p*K*_a_ counts into account, we identified a shortlist of consistently well-performing methods for macroscopic p*K*_a_ predictions, shown in Table 2. The criteria for selection were the overall ranking in Top 10 according to RMSE, MAE, R^2^, and Kendall’s Tau and also having a combined unmatched p*K*_a_ (extra and missing p*K*_a_s) count less than 8 (a third of the number of compounds). We ranked methods in ascending order for RMSE and MAE and in descending order for R^2^, and Kendall’s Tau to determine methods. Then, we took the intersection set of Top 10 methods according to each statistic to determine the consistently-well performing methods. This resulted in a list of four methods that are consistently well-performing across all criteria.

Consistently well-performing methods for macroscopic p*K*_a_ prediction included methods from all categories. Two methods in the QM+LEC category were *xvxzd* (DSD-BLYP-D3(BJ)/def2-TZVPD//PBEh-3c[DCOSMO-RS] + RRHO(GFN-xTB[GBSA]) +Gsolv(COSMO-RS[TZVPD]) and linear fit) and (8xt50) (ReSCoSS conformations // DSD-BLYP-D3 reranking // COSMOtherm pKa) and both used COSMO-RS. Empirical p*K*_a_ predictions with top performance were both proprietary software. From QSPR and LFER categories, *gyuhx* (Simulations Plus) and *xmymhm* (ACD/pKa Classic) were consistently well-performing methods. The Simulation Plus p*K*_a_ prediction method consisted of 10 artificial neural network ensembles trained on 16,000 compounds for 10 classes of ionizable atoms, with the ionization class of each atom determined using an assigned atom type and local molecular environment [48]. The ACD/pKa Classic method was trained on 17,000 compounds, uses Hammett-type equations, and captures effects related to tautomeric equilibria, covalent hydration, resonance effects, and *α, β*-unsaturated systems [38].

Figure 4 plots predicted vs. experimental macroscopic p*K*_a_ predictions of four consistently well-performing methods, a representative average method, and the null method(*5nm4j*). We selected the method with the highest RMSE below the median of all methods as the representative method with average performance: *2ii2g* (EC-RISM/MP2/cc-pVTZ-P2-q-noThiols-2par).

**Figure 4.**
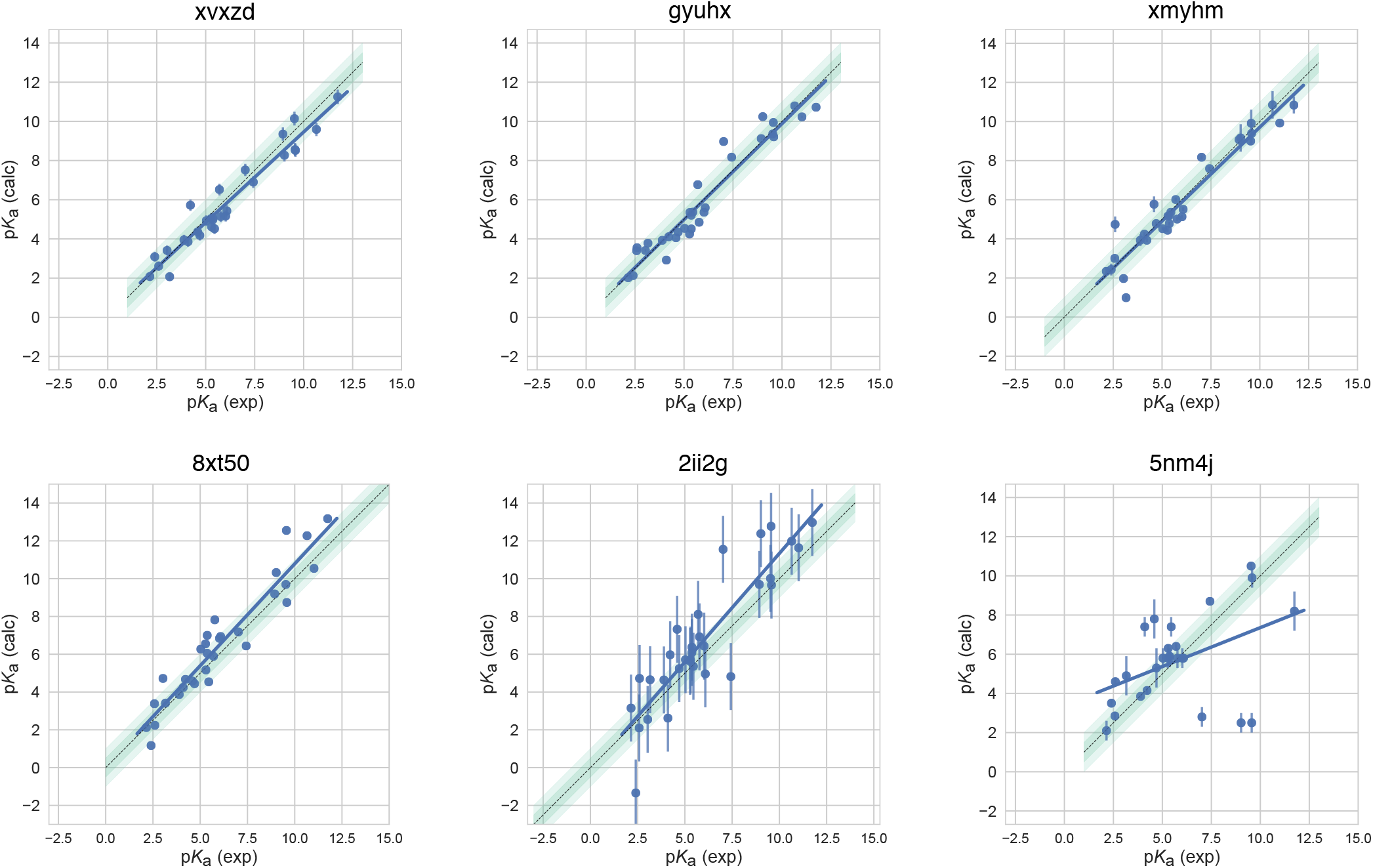
Predicted vs. experimental macroscopic p*K*_a_ prediction for four consistently well-performing methods, a representative method with average performance (*2ii2g*), and the null method (*5nm4j*). When submissions were ranked according to RMSE, MAE, R^2^, and *τ*, four methods ranked in the Top 10 consistently in each of these metrics. Dark and light green shaded areas indicate 0.5 and 1.0 units of error. Error bars indicate standard error of the mean of predicted and experimental values. Experimental p*K*_a_ SEM values are too small to be seen under the data points. EC-RISM/MP2/cc-pVTZ-P2-q-noThiols-2par method (*2ii2g*) was selected as the representative method with average performance because it is the method with the highest RMSE below the median.

#### 3.1.2 Which chemical properties are driving macroscopic p*K*_a_ prediction failures?

In addition to comparing the performance of methods that participated in the SAMPL6 Challenge, we also wanted to analyze macroscopic p*K*_a_ predictions from the perspective of challenge molecules and determine whether particular compounds suffer from larger inaccuracy in p*K*_a_ predictions. The goal of this analysis is to provide insighton which molecular properties or moieties might be causing larger p*K*_a_ prediction errors. In Fig. 5, 2D depictions of the challenge molecules are presented with MAE calculated fortheir macroscopic p*K*_a_ predictions over all methods, based on Hungarian match. For multiprotic molecules, the MAE was averaged over all the p*K*_a_ values. For the analysis of p*K*_a_ prediction accuracy observed for each molecule, MAE is a more appropriate statistical value than RMSE for following global trends, as it is less sensitive to outliers than the RMSE.

**Figure 5.**
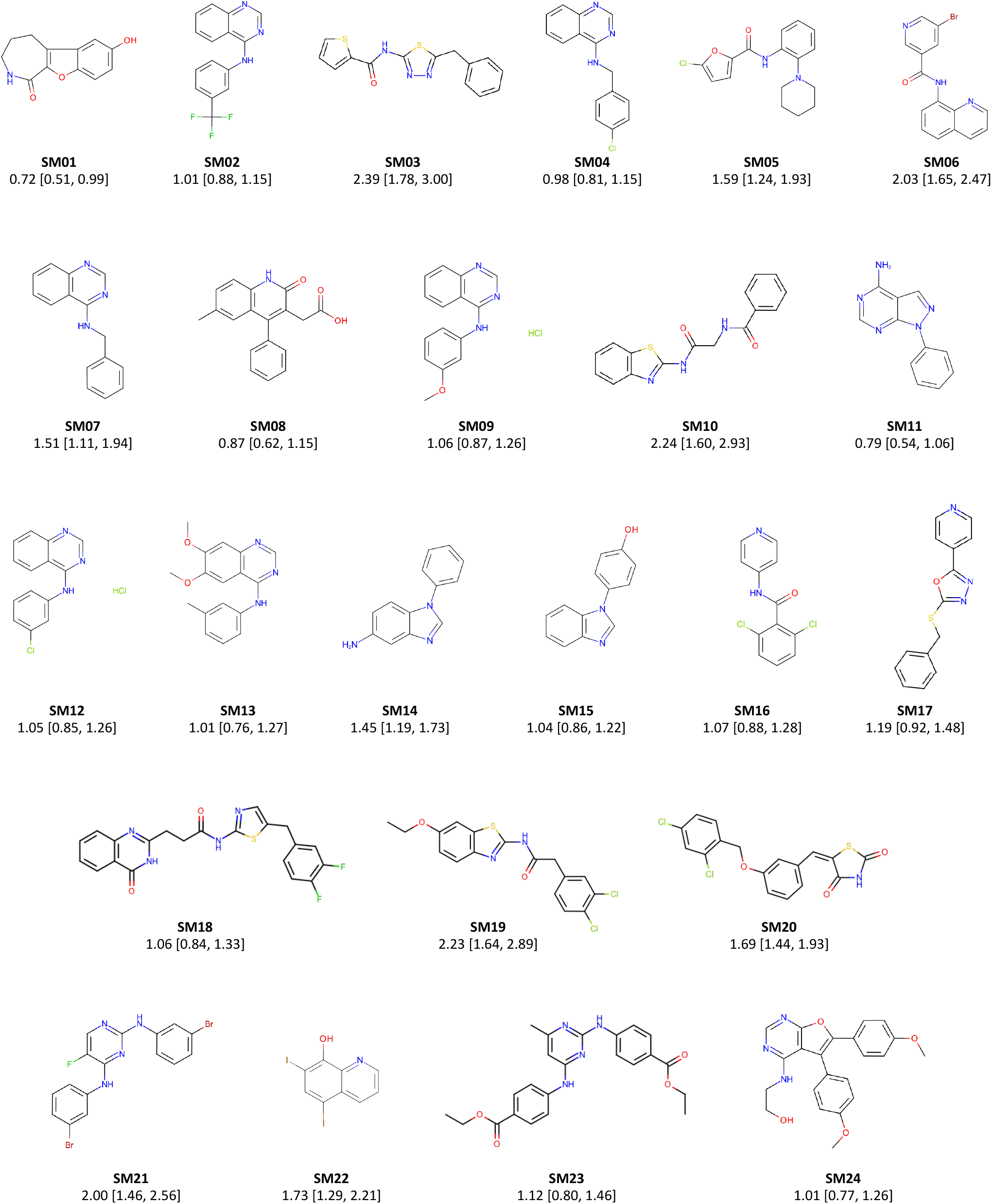
Molecules from the SAMPL6 Challenge with MAE calculated for all macroscopic p*K*_a_ predictions. The MAE calculated over all prediction methods indicates which molecules had the lowest prediction accuracy in the SAMPL6 Challenge. MAE values calculated for each molecule include all the matched p*K*_a_ values. SM06, SM14, SM15, SM16, SM18, and SM22 were multiprotic. Hungarian matching algorithm was employed for pairing experimental and predicted p*K*_a_ values. MAE values are reported with 95% confidence intervals.

A comparison of the prediction accuracy of individual molecules is shown in Fig. 6. In Fig. 6A, the MAE for each molecule is shown considering all blind predictions and reference calculations. A cluster of molecules marked orange and red have higher than average MAE. Molecules marked red (SM06, SM21, and SM22) are the only compounds in the SAMPL6 dataset with bromo or iodo groups and they suffered a macroscopic p*K*_a_ prediction error in the range of 1.7-2.0 p*K*_a_ units in terms of MAE. Molecules marked orange (SM03, SM10, SM18, SM19, and SM20) have sulfur-containing heterocycles, and all these molecules except SM18 have MAE larger than 1.6 p*K*_a_ units. Despite containing a thiazole group, SM18 has a low prediction MAE. SM18 is the only compound with three experimental p*K*_a_ values, and we suspect the presence of multiple experimental p*K*_a_ values could have a masking effect on the errors captured by the MAE when the Hungarian matching scheme is used due to more potential pairing choices that may artificially lower the error.

**Figure 6.**
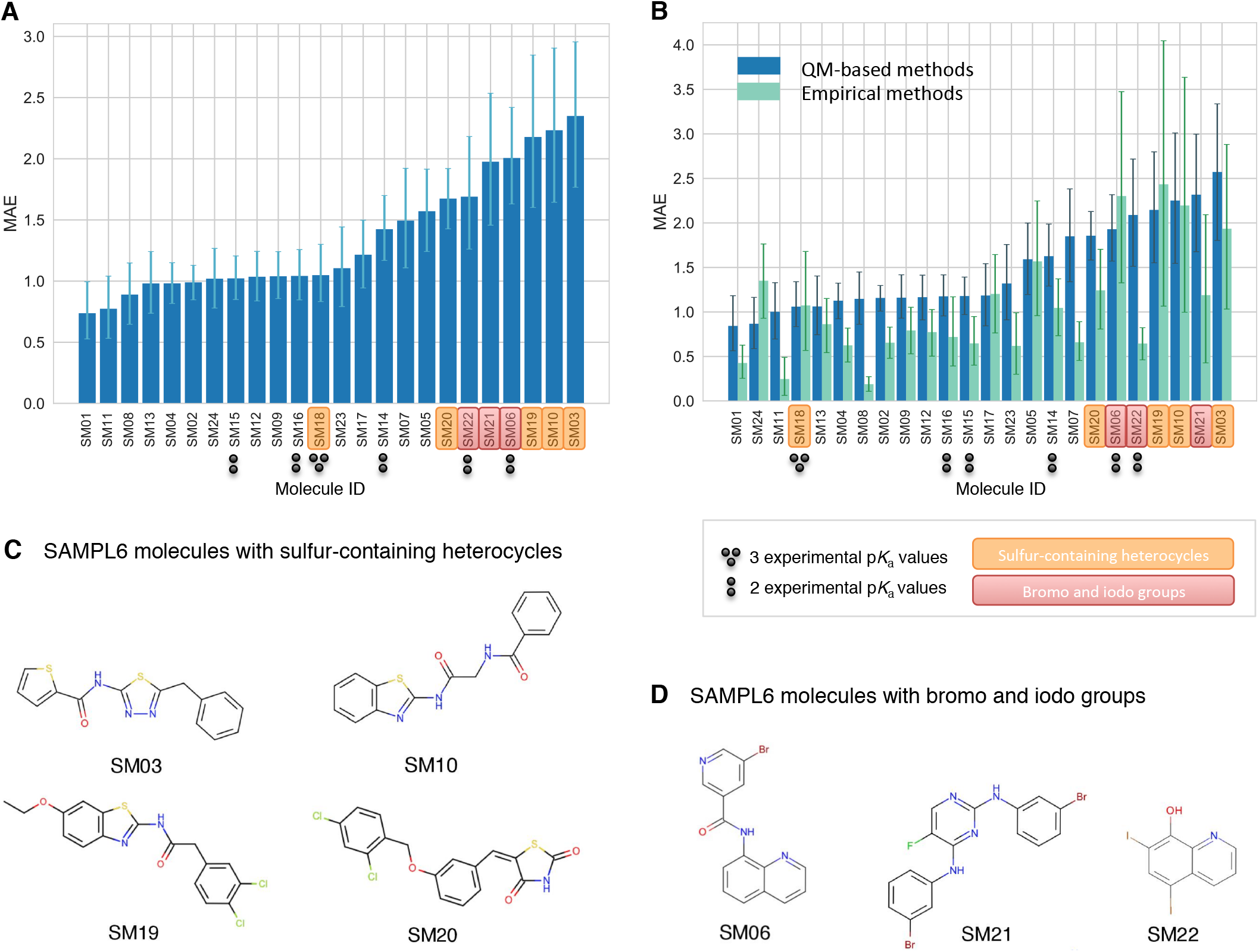
Average prediction accuracy calculated over all prediction methods was poorer for molecules with sulfur-containing heterocycles, bromo, and iodo groups. **(A)** MAE calculated for each molecule as an average of all methods. **(B)** MAE of each molecule broken out by method category. QM-based methods (blue) include QM predictions with or without linear empirical correction. Empirical methods (green) include QSAR, ML, DL, and LFER approaches. **(C)** Depiction of SAMPL6 molecules with sulfur-containing heterocycles. **(D)** Depiction of SAMPL6 molecules with iodo and bromo groups.

We separately analyzed the MAE of each molecule for empirical (LFER and QSPR/ML) and QM-based physical methods (QM, QM+LEC, and QM+MM) to gain additional insight into prediction errors. Fig. 6B shows that the difficulty of predicting p*K*_a_ values of the same subset of molecules was a trend conserved in the performance of physical methods. For QM-based methods, sulfur-containing heterocycles, amides proximal to aromatic heterocycles, and compounds with iodo and bromo substitutions have lower p*K*_a_ prediction accuracy.

The SAMPL6 p*K*_a_ set consists of only 24 small molecules and lacks multiple examples of many moieties, limiting our ability to determine with statistical significance which chemical substructures cause greater errors in p*K*_a_ predictions. Still, the trends observed in this challenge point to molecules with iodo-, bromo-, and sulfur-containing heterocycles as having systematically larger prediction errors in macroscopic p*K*_a_ value. We hope that reporting this observation will lead to the improvement of methods for similar compounds with such moieties.

We have also looked for correlation with molecular descriptors for finding other potential explanations as to why macroscopic p*K*_a_ prediction errors were larger for certain molecules. While testing the correlation between errors and many molecular descriptors, it is important to account for the possibility of spurious correlations. We haven’t observed any statistically significant correlation between numerical p*K*_a_ predictions and the descriptors we have tested. First, having more experimental p*K*_a_ values (Fig. 6A) did not seem to be associated with poorer p*K*_a_ prediction performance. Still, we need to keep in mind that multiprotic compounds were sparsely represented in the SAMPL6 set (5 molecules with 2 macroscopic p*K*_a_ values and one with 3 macroscopic p*K*_a_). Second, we checked the following other descriptors: presence of an amide group, molecular weight, heavy atom count, rotatable bond count, heteroatom count, heteroatom-to-carbon ratio, ring system count, maximum ring size, and the number of microstates (as enumerated for the challenge). Correlation plots and R^2^ values can be seen in Fig. S2.

We had suspected that p*K*_a_ prediction methods may perform better for moderate values (4-10) than extreme values as molecules with extreme p*K*_a_ values are less likely to change ionization states close to physiological pH. To test this we look at the distribution of absolute errors calculated for all molecules and challenge predictions binned by experimental p*K*_a_ value 2 p*K*_a_ unit increments. As can be seen in Fig. S3B, the value of true macroscopic p*K*_a_ values was not a factor affecting the prediction error seen in SAMPL6 Challenge.

Fig. 7B is helpful to answer the question “Are there molecules with consistently overestimated or underestimated p*K*_a_ values?”. This ridge plots show the error distribution of each experimental p*K*_a_. SM02_pKa1, SM04_pKa1, SM14_pKa1, and SM21_pKa1 were underestimated, predicting lower protein affinity by more than 1 p*K*_a_ unit by majority of the prediction methods. SM03_pKa1, SM06_pKa2, SM19_pKa1, and SM20_pKa1 were overestimated by the majority of the prediction methods by more than 1 p*K*_a_ unit. SM03_pKa1, SM06_pKa2, SM10_pKa1, SM19_pKa1, and SM22_pKa1 have the highest spread of errors and were less accurately predicted overall.

**Figure 7.**
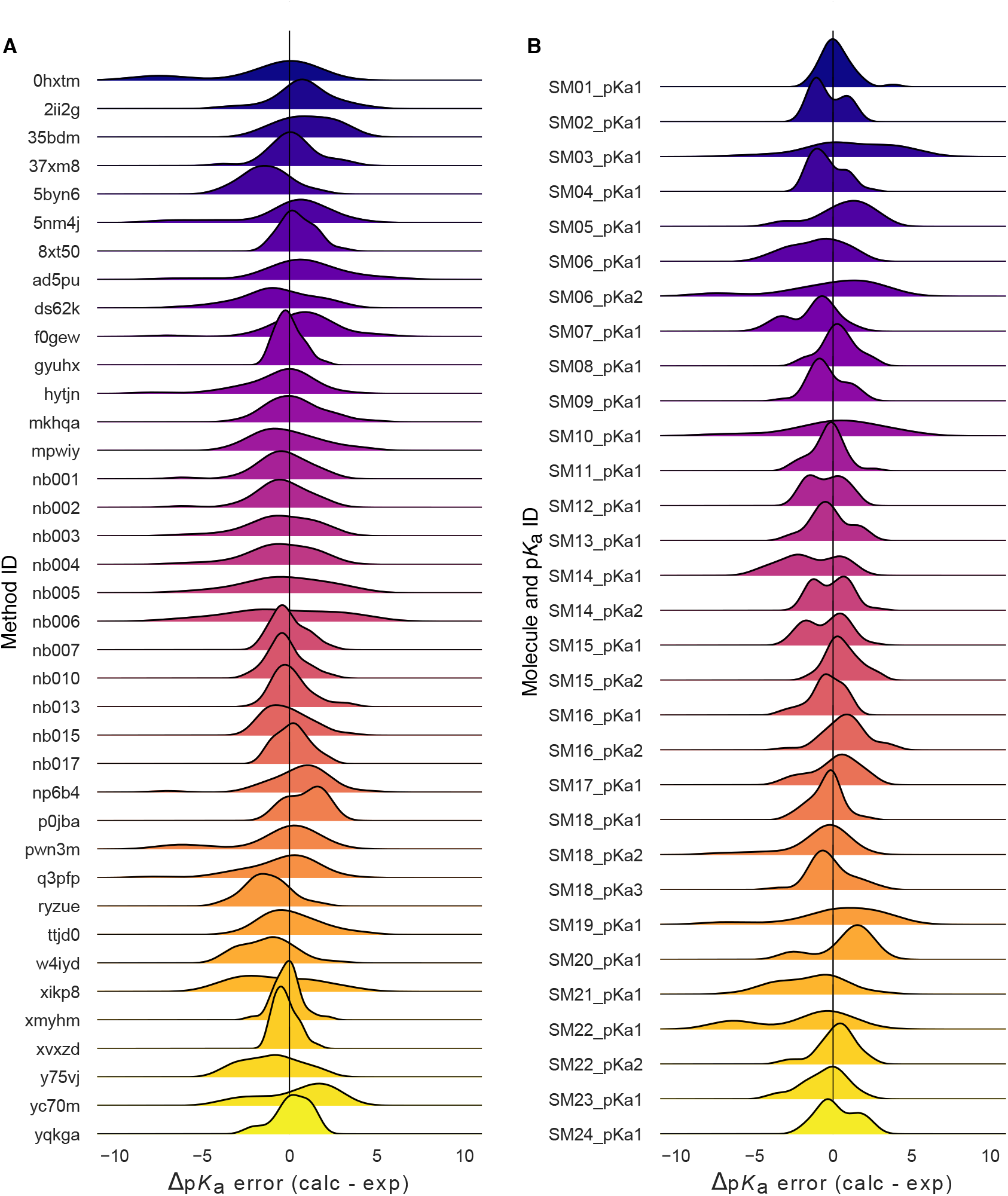
Macroscopic p*K*_a_ prediction error distribution plots show how prediction accuracy varies across methods and individual molecules. **(A)** p*K*_a_ prediction error distribution for each submission for all molecules according to Hungarian matching. **(B)** Error distribution for each SAMPL6 molecule for all prediction methods according to Hungarian matching. For multiprotic molecules, p*K*_a_ ID numbers (pKa1, pKa2, and pKa3) were assigned in the direction of increasing experimental p*K*_a_ value.

### 3.2 Analysis of microscopic p*K*_a_ predictions using microstates determined by NMRfor 8 molecules

The most common approach for analyzing microscopic p*K*_a_ prediction accuracy has been to compare it to experimental macroscopic p*K*_a_ data, assuming experimental p*K*_a_ values describe titrations of distinguishable sites and, therefore, correspond to microscopic p*K*_a_s. But this typical approach fails to evaluate methods at the microscopic level.

Analysis of microscopic p*K*_a_ predictions for the SAMPL6 Challenge was not straightforward due to the lack of experimental data with microscopic resolution of the titratable sites and their associated microscopic p*K*_a_s. For 24 molecules, macroscopic p*K*_a_ values were determined with the spectrophotometric method. For 18 molecules, a single macroscopic titration was observed, and for 6 molecules multiple experimental p*K*_a_ values were observed and characterized. For 18 molecules with a single experimental p*K*_a_, it is probable that the molecules are monoprotic and, therefore, macroscopic p*K*_a_ value is equal to the microscopic p*K*_a_. There is, however, no direct experimental evidence supporting this hypothesis aside from the support from computational predictions, such as the predictions by ACD/pKa Classic. There is always the possibility that the macroscopic p*K*_a_ observed is the result of two different titrations overlapping closely with respect to pH if any charge state has more than one tautomer. We did not want to bias the blind challenge analysis with any prediction method. Therefore, we believe analyzing the microscopic p*K*_a_ predictions via Hungarian matching to experimental values with the assumption that the 18 molecules have a single titratable site is not the best approach. Instead, an analysis at the level of macroscopic p*K*_a_ values is much more appropriate when a numerical matching scheme is the only option to evaluate predictions using macroscopic experimental data.

For a subset of eight molecules, dominant microstates were inferred from NMR experiments. Six of these molecules were monoprotic and two were multiprotic. This dataset was extremely useful for guiding the assignment between experimental and predicted p*K*_a_ values based on microstates. In this section, we present the performance evaluations of microscopic p*K*_a_ predictions for only the 8 compounds with experimentally-determined dominant microstates.

#### 3.2.1 Microstate-based matching revealed errors masked by p*K*_a_ value-based matching between experimental and predicted p*K*_a_s

Comparing microscopic p*K*_a_ predictions directly to macroscopic experimental p*K*_a_ values with numerical matching can lead to underestimation of errors. To demonstrate how numerical matching often masks p*K*_a_ prediction errors, we compared the performance analysis done by Hungarian matching to that from microstate-based matching for 8 molecules presented in Fig. 8A. RMSE calculated for microscopic p*K*_a_ predictions matched to experimental values via Hungarian matching is shown in Fig. 8B, while Fig. 8C shows RMSE calculated via microstate-based matching. The Hungarian matching incorrectly leads to significantly (and artificially) lower RMSE compared to microstate-based matching. The reason is that the Hungarian matching assigns experimental p*K*_a_ values to predicted p*K*_a_ values only based on the closeness of the numerical values, without consideration of the relative population of microstates and microstate identities. Because of this, a microscopic p*K*_a_ value that describes a transition between very low population microstates (high energy tautomers) can be assigned to the experimental p*K*_a_ if it has the closest p*K*_a_ value. This is not helpful because, in reality, the microscopic p*K*_a_ values that influence the observable macroscopic p*K*_a_ the most are the ones with higher microstate populations (transitions between low energy tautomers).

**Figure 8.**
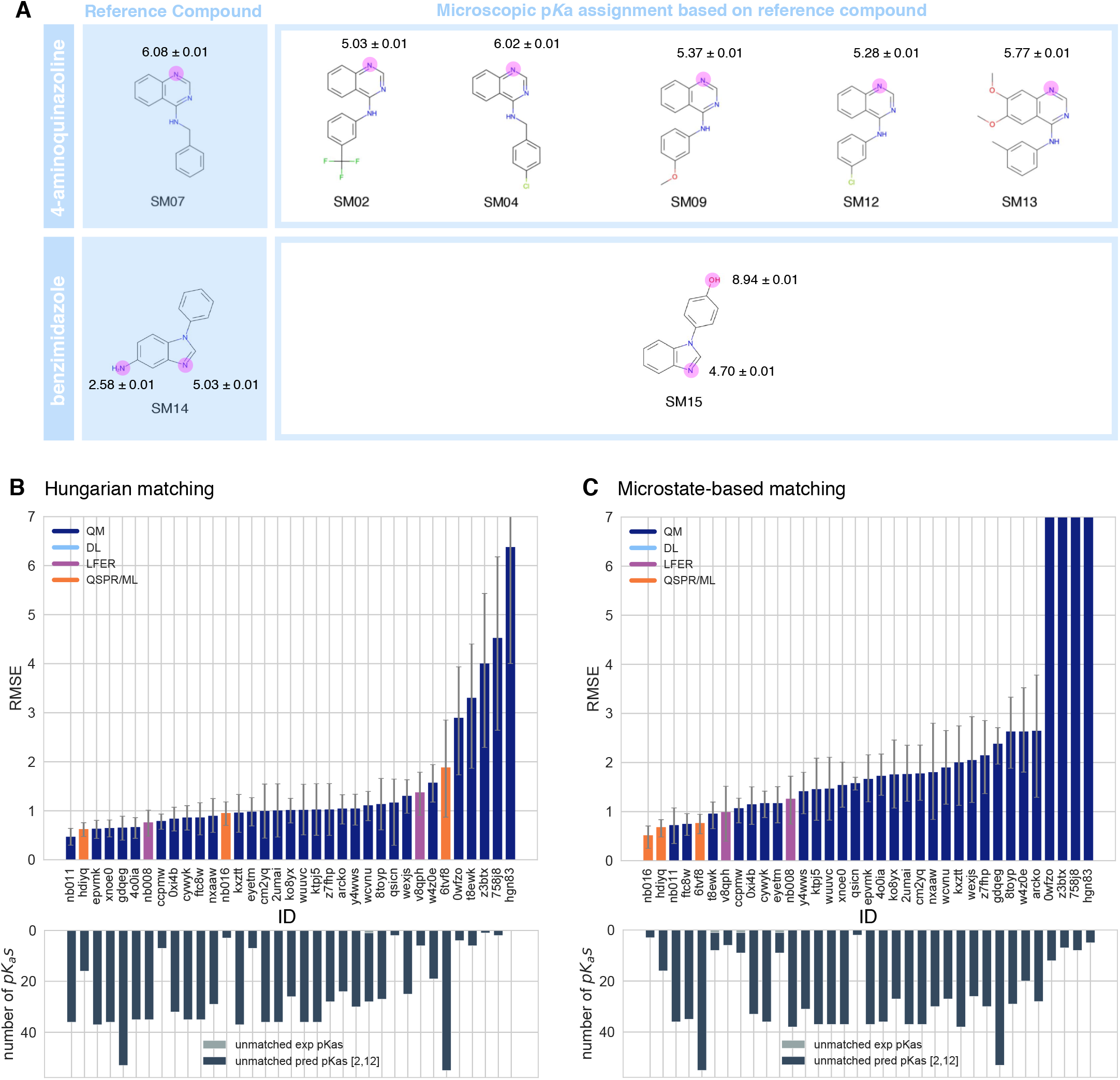
NMR determination of dominant microstates allowed in-depth evaluation of microscopic p*K*_a_ predictions for 8 compounds. **A** Dominant microstate sequence of two compounds (SM07 and SM14) were determined by NMR [8]. Based on these reference compounds, the dominant microstates of 6 related compounds were inferred and experimental p*K*_a_ values were assigned to titratable groups with the assumption that only the dominant microstates have significant contributions to the experimentally observed p*K*_a_. **B** RMSE vs. submission ID and unmatched p*K*_a_ vs. submission ID plots for the evaluation of microscopic p*K*_a_ predictions of 8 molecules by Hungarian matching to experimental macroscopic p*K*_a_ values. **C** RMSE vs. submission ID and unmatched p*K*_a_ vs. submission ID plots showing the evaluation of microscopic p*K*_a_ predictions of 8 molecules by microstate-based matching between predicted microscopic p*K*_a_s and experimental macroscopic p*K*_a_ values. Submissions *0wfzo, z3btx, 758j8*, and *hgn83* have RMSE values bigger than 10 p*K*_a_ units which are beyond the y-axis limits of subplot **C** and **B**. RMSE is shown with error bars denoting 95% confidence intervals obtained by bootstrapping over the challenge molecules. Lower bar plots show the number of unmatched experimental p*K*_a_s (light grey, missing predictions) and the number of unmatched p*K*_a_ predictions (dark grey, extra predictions) for each method between pH 2 and 12. Submission IDs are summarized in Table 1.

The number of unmatched predicted microscopic p*K*_a_s is shown in the lower bar plots of Fig. 8B and C, to emphasize the large number of microscopic p*K*_a_ predictions submitted by many methods. In the case of microscopic p*K*_a_, the number of unmatched predictions does not indicate an error in the form of an extra predicted p*K*_a_, because the spectrophotometric experiments do not capture all microscopic p*K*_a_s theoretically possible (transitions between all pairs of microstates that differ by one proton). p*K*_a_s of transitions to and from very high energy tautomers are very hard to measure by experimental methods, including the most sensitive methods like NMR. Prediction of extra microscopic p*K*_a_ values can cause underestimation of prediction errors when numerical matching algorithms such as Hungarian matching are used. We also checked how often Hungarian matching led to the correct matches between predicted and experimental p*K*_a_ in terms of the microstate pairs, i.e., how often the microstate pair of the Hungarian match recapitulates the dominant microstate pair of the experiment. The overall accuracy of microstate pair matching was found to be low for the SAMPL6 Challenge submission. Fig. S4 shows that for most methods the predicted microstate pair selected by the Hungarian match did not correspond to the experimentally-determined microstate pair. This means lower RMSE (better accuracy) performance statistics obtained from Hungarian matching are artificially low. This problem could be avoided by matching experimental and predicted values on the basis of microstate IDs, if experimental microscopic assignments are available.

Unfortunately, we were only able to perform this more reliable microstate-based analysis for a subset of compounds. The conclusions in this section reflect only eight compounds with limited structural diversity: Six molecules with 4-aminoquinazoline and two with benzimidazole scaffolds, with a total of 10 p*K*_a_ values. The sequences of dominant microstates for SM07 and SM14 were determined by NMR experiments directly [8], while dominant microstates of their derivatives were inferred by taking them as a reference (Fig. 8). Although we believe that microstate-based evaluation is more informative, the lack of a large experimental dataset limits the conclusions to a very narrow chemical diversity. Still, microstate-based matching revealed errors masked by p*K*_a_ value-based matching between experimental and predicted p*K*_a_s.

#### 3.2.2 Accuracy of p*K*_a_ predictions evaluated by microstate-based matching

Both accuracy- and correlation-based statistics were calculated for the predicted microscopic p*K*_a_ values after microstate-based matching. RMSE, MAE, ME, R^2^, and Kendall’s Tau results of each method are shown in Fig. 8C and Fig. 9. Atable of the calculated statistics can be found in Table S4. Due to the small number of data points in this set, correlation-based statistics have large uncertainties and thus have less utility for distinguishing better-performing methods. Therefore, we focused more on accuracy-based metrics for the analysis of microscopic p*K*_a_s than correlation-based metrics. In terms of accuracy of predicted microscopic p*K*_a_ values, all three QSPR/ML based methods (*nb016* (MoKa), *hdiyq* (Simulations Plus), *6tvf8* (OE Gaussian Process)), three QM-based methods (*nb011* (Jaguar), *ftc8w* (EC-RISM/MP2/cc-pVTZ-P2-q-noThiols-2par), *t8ewk* (COSMOlogic_FINE17)), and one LFER method (*v8qph* (ACD/pKa GALAS)) achieved RMSE lower than 1 p*K*_a_ unit. The same six methods also have the lowest MAE.

**Figure 9.**
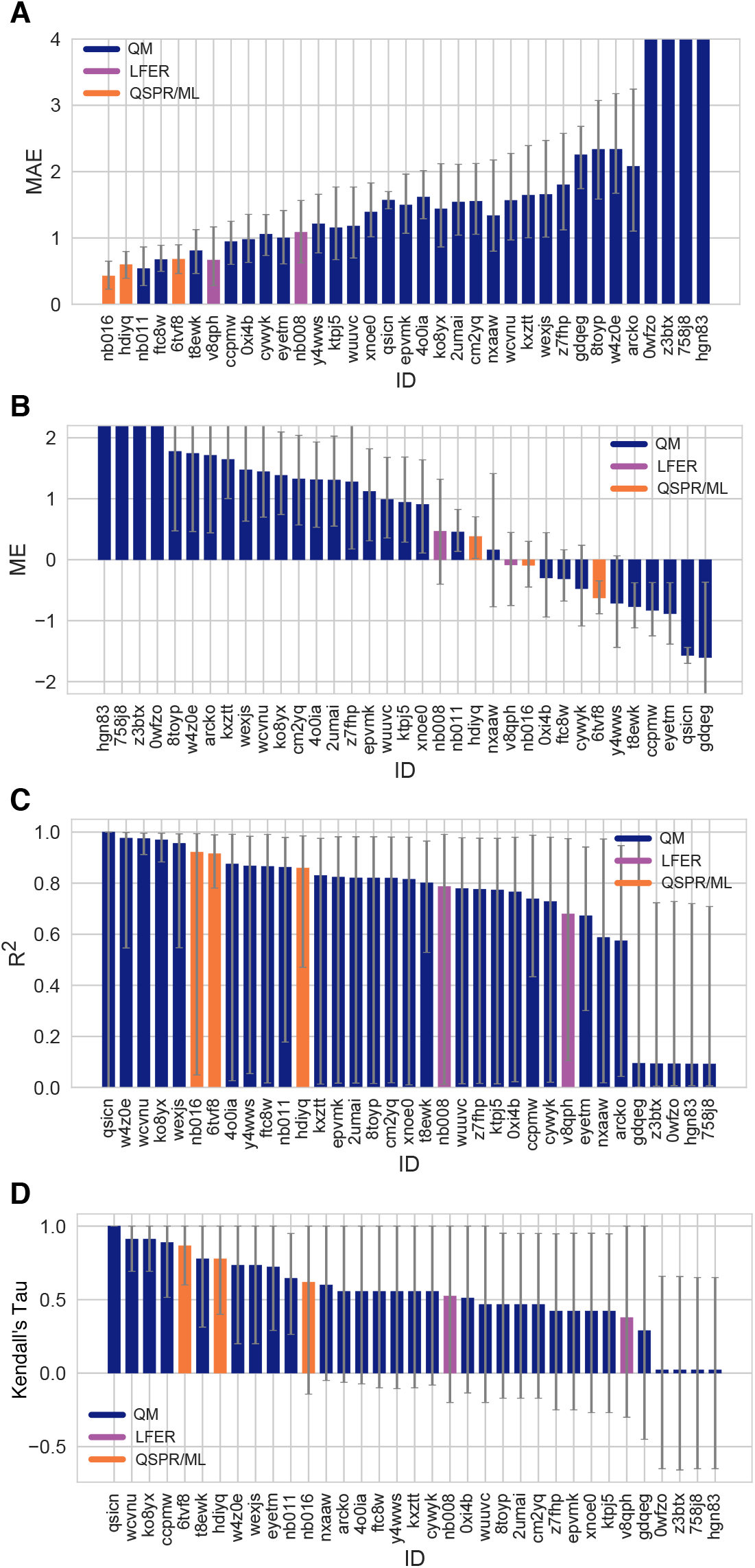
Additional performance statistics for microscopic p*K*_a_ predictions for 8 molecules with experimentally determined dominant microstates. Microstate-based matching was performed between experimental p*K*_a_ values and predicted microscopic p*K*_a_ values. Mean absolute error (MAE), mean error (ME), Pearson’s R^2^, and Kendall’s Rank Correlation Coefficient Tau (***τ***) are shown, with error bars denoting 95% confidence intervals obtained by bootstrapping over challenge molecules. Methods are indicated by their submission IDs. Submissions are colored by their method categories. Refer to Table 1 for submission IDs and method names. Submissions *0wfzo, z3btx, 758j8*, and *hgn83* have MAE and ME values bigger than 10 p*K*_a_ units which are beyond they-axis limits of subplots **A** and **B**. A large number and wide variety of methods have statistically indistinguishable performance based on correlation statistics (**C** and **D**), in part because of the relatively small dynamic range and small size of the set of 8 molecules.

#### 3.2.3 Evaluation of dominant microstate prediction accuracy

For many computational chemistry approaches, including structure-based modeling of protein-ligand interactions, predicting the ionization state and the exact position of protons is necessary to establish what to include in the modeled system. In addition to being able to predict p*K*_a_ values accurately, we require p*K*_a_ prediction methods to be able to capture microscopic protonation states accurately. Even when the predicted p*K*_a_ value is accurate, the predicted protonation sites can be incorrect, leading to potentially large modeling errors in quantities such as the computed free energy of binding. Therefore, we assessed whether methods participating in the SAMPL6 p*K*_a_ Challenge were correctly predicting the sequence of dominant microstates, i.e., dominant tautomers of each charge state observed between pH 2 and 12.

Fig. 10 shows how well methods perform for predicting the dominant microstate, as analyzed for eight compounds with available experimental microstate assignments. The dominant microstate sequence is essentially the sequence of states that are most visible experimentally due to their higher fractional population and relative free energy within the tautomers at each charge. To extract the dominant tautomers predicted for the sequence of ionization states of each method, the relative free energy of microstates were first calculated at reference pH 0 [26]. To subsequently determine the dominant microstate at each formal charge, we selected the lowest energy tautomer for each ionization state based on the relative microstate free energies calculated at pH 0. The choice of reference pH is arbitrary, as relative free energy difference between tautomers of the same charge is always constant with respect to pH. This analysis was performed only for the charges −1,0,1, and 2—the charge range captured by NMR experiments. Predicted and experimental dominant microstates were then compared for each charge state to calculate the fraction of correctly predicted dominant tautomers. This value is reported as the *dominant microstate accuracy* for all charge states (Fig. 10A).

**Figure 10.**
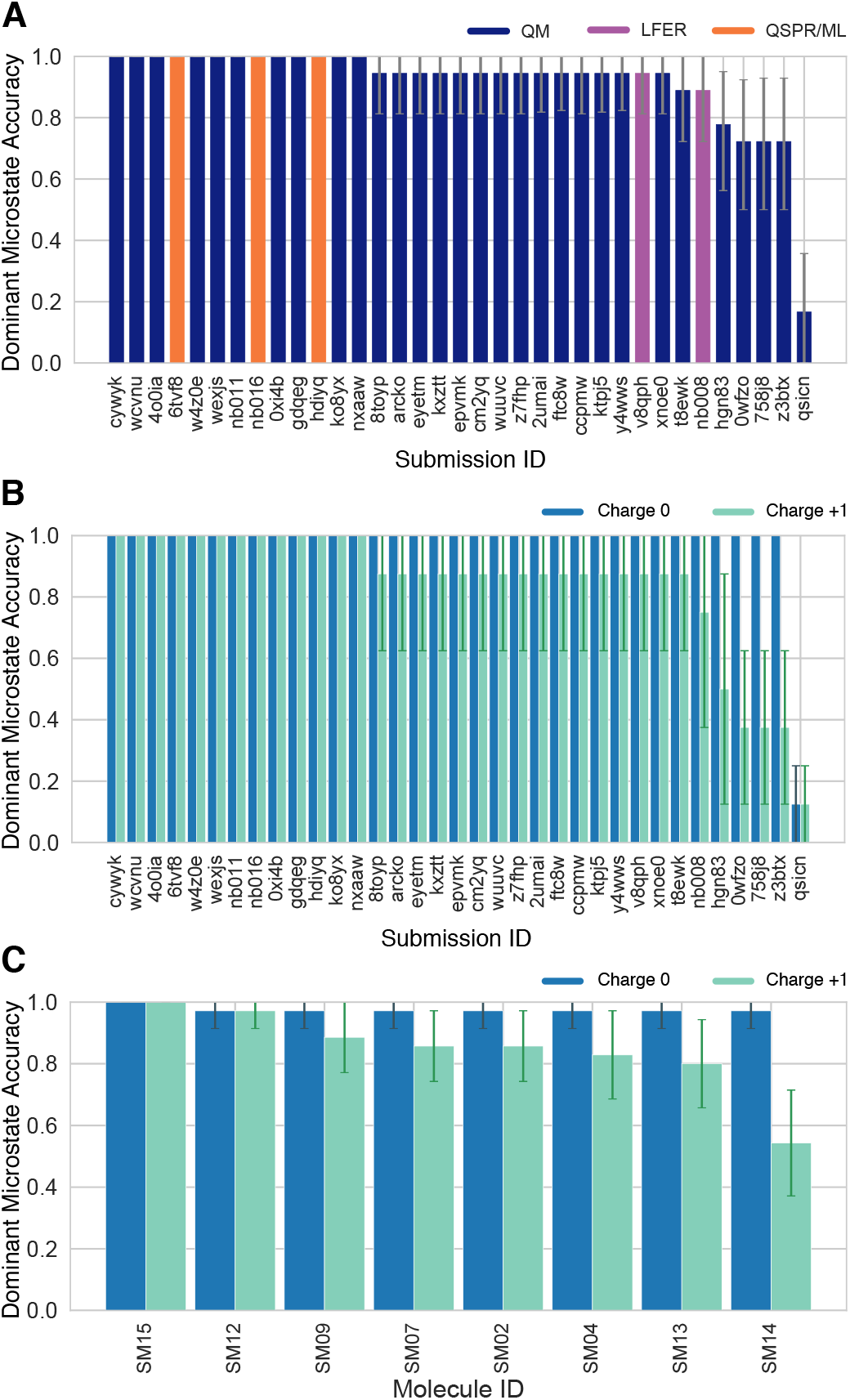
Some methods predicted the sequence of dominant tautomers inaccurately. Prediction accuracy of the dominant microstate of each charged state was calculated using the dominant microstate sequence determined by NMR for 8 molecules as reference. **(A)** Dominant microstate accuracy vs. submission ID plot was calculated considering all the dominant microstates seen in the experimental microstate dataset of 8 molecules. **(B)** Dominant microstate accuracy vs. submission ID plot was generating considering only the dominant microstates of charge 0 and +1 seen in the 8 molecule dataset. The accuracy of each molecule is broken out by the total charge of the microstate. **(C)** Dominant microstate prediction accuracy calculated for each molecule averaged over all methods. In **(B)** and **(C)**, the accuracy of predicting the dominant neutral tautomer is shown in blue and the accuracy of predicting the dominant +1 charged tautomer is shown in green. Error bars denoting 95% confidence intervals obtained by bootstrapping.

Many of the methods which participated in the challenge made errors in predicting the dominant microstate. 10 QM and 3 QSPR/ML methods did not make any mistakes in dominant microstate predictions, although, they are expected to make mistakes in the relative population of tautomers (free energy difference between microstates) as reflected by the p*K*_a_ value errors. While all participating QSPR/ML methods showed good performance in dominant microstate prediction, LFER and some QM methods made mistakes. The accuracy of the predicted dominant neutral tautomers was perfect for all methods, except *qsicn* (Fig. 10B), but errors in predicting the major tautomer of charge +1 were much more frequent. 22 out of 35 prediction sets made at least one error in predicting the lowest energy tautomer with +1 charge. We didn’t include ionization states with charges −1 and +2 in this assessment because we had only one compound with these charges in the dataset. Nevertheless, errors in predicting the dominant tautomers seem to be a bigger problem for charged tautomers than the neutral tautomer.

Only eight compounds had data on the sequence of dominant microstates. Therefore conclusions on the performance of methods in terms of dominant tautomer prediction are limited to this limited chemical diversity (benzimidazole and 4-aminoquinazoline derivatives). We present this analysis as a prototype of how microscopic p*K*_a_ predictions should be evaluated. Hopefully, future evaluations can be performed with larger experimental datasets following the strategy we demonstrated here in order to reach broad conclusions about which methods are better for capturing dominant microstates and ratios of tautomers. Even if experimental microscopic p*K*_a_ measurement data is not available, experimental dominant tautomer determinations are still informative for assessing computational predictions.

The most frequent misprediction was the major tautomer of the SM14 cationic form, as shown in Fig. 10. This figure shows the accuracy of the predicted dominant microstate calculated for individual molecules and for charge states 0 and +1, averaged over all prediction methods. SM14, the molecule that exhibits the most frequent error in the predicted dominant microstate, has two experimental p*K*_a_ values that were 2.4 p*K*_a_ units apart, and we suspect that could be a contributor to the difficulty of predicting microstates accurately. Other molecules are monoprotic (4-aminoquinazolines) or their experimental p*K*_a_ values are very well separated (SM14,4.2 p*K*_a_ units). It would be very interesting to expand this assessment to a larger variety of drug-like molecules to discover for which structures tautomer predictions are more accurate and for which structures computational predictions are not as reliable.

#### 3.2.4 Consistently well-performing methods for microscopic p*K*_a_ predictions

We have identified different criteria for determining consistently top-performing predictions of microscopic p*K*_a_ than macroscopic p*K*_a_: having perfect dominant microstate prediction accuracy, unmatched p*K*_a_ count of 0, and ranking in the top 10 according to RMSE and MAE. Correlation statistics were not found to have utility for discriminating performance due to large uncertainties in these statistics for a small dataset of 10 p*K*_a_ values. Unmatched predicted p*K*_a_ count was also not considered since experimental data was only informative for the p*K*_a_ between dominant microstates and did not capture all the possible theoretical transitions between microstate pairs. Table 3 reports six methods that have consistent good performance according to many metrics, although evaluated only for the 8 molecule set due to limitations of the experimental dataset. Six methods were divided evenly between methods of QSPR/ML category and QM category. *nb016* (MoKa), *hdiyq* (Simulations Plus), and *6tvf8* (OE Gaussian Process) were QSPR and ML methods that performed well. *nb011* (Jaguar), *0xi4b*(EC-RISM/B3LYP/6-311+G(d,p)-P2-phi-noThiols-2par), and *cywyk* (EC-RISM/B3LYP/6-311+G(d,p)-P2-phi-noThiols-2par) were QM predictions with linear empirical corrections with good performance with microscopic p*K*_a_ predictions.

The Simulations Plus p*K*_a_ prediction method is the only method that appeared to be consistently well-performing in both the assessment for macroscopic and microscopic p*K*_a_ prediction (*gyuhx* and *hdiyq*). However, it is worth noting that two methods that were in the list of consistently top-performing methods for macroscopic p*K*_a_ predictions lacked equivalent submissions of their underlying microscopic p*K*_a_ predictions, and therefore could not be evaluated at the microstate level. These methods were *xmyhm* (ACD/pKa Classic) and *xvxzd*(DSD-BLYP-D3(BJ)/def2-TZVPD//PBEh-3c[DCOSMO-RS] + RRHO(GFN-xTB[GBSA]) + Gsolv(COSMO-RS[TZVPD]) and linear fit).

### 3.3 How do p*K*_a_ prediction errors impact protein-ligand binding affinity predictions?

p*K*_a_ predictions provide a key input for computational modeling of protein-ligand binding with physical methods. The SAMPL6 p*K*_a_ Challenge focused only on small molecule p*K*_a_ prediction and showed how p*K*_a_ prediction accuracy observed can impact the modeling of ligands. Many affinity prediction methods such as docking, MM/PBSA, MM/GBSA, absolute or alchemical relative free energy calculation methods predict the affinity of the ligand to a receptor using a fixed protonation state for both ligand and receptor. These models can sensitively depend upon p*K*_a_ and dominant tautomer predictions for determining possible protonation states of the ligand in the aqueous environment and in a protein complex, as well as the free energy penalty to access those states [4]. The accuracy of p*K*_a_ predictions can become a limitation for the performance of physical models that try to quantitatively describe molecular association.

In terms of ligand protonation states, there are two ways in which p*K*_a_ prediction errors can influence the prediction accuracy for protein-ligand binding free energies as depicted in Fig. 11. The first scenario is when a ligand is present in aqueous solution in multiple protonation states (Fig. 11A). When only the minor aqueous protonation state contributes to protein-ligand complex formation, the overall binding free energy (Δ*G_bind_*) needs to be calculated as the sum of binding free energy of the minor state and the protonation penalty of that state (Δ*G_prot_*). Δ*G_prot_* is a function of both pH and p*K*_a_. A1 unit of error in predicted p*K*_a_ would lead to 1.36 kcal/mol error in overall binding free energy if the protonation state with the minor population binds the protein and this minor protonation state is *correctly* selected to model the free energy of binding; if the incorrect dominant protonation state for the complex is selected, the dominant contribution to the free energy of binding may be missed entirely, leading to much larger modeling errors in the binding free energy. Other scenarios—in which multiple protonation states can be significantly populated in complex—can lead to more complex scenarios in which the errors in predicted p*K*_a_ propagate in more complex ways. The equations in Fig. 11A show the overall free energy for a simple thermodynamic cycle involving multiple protonation states.

**Figure 11.**
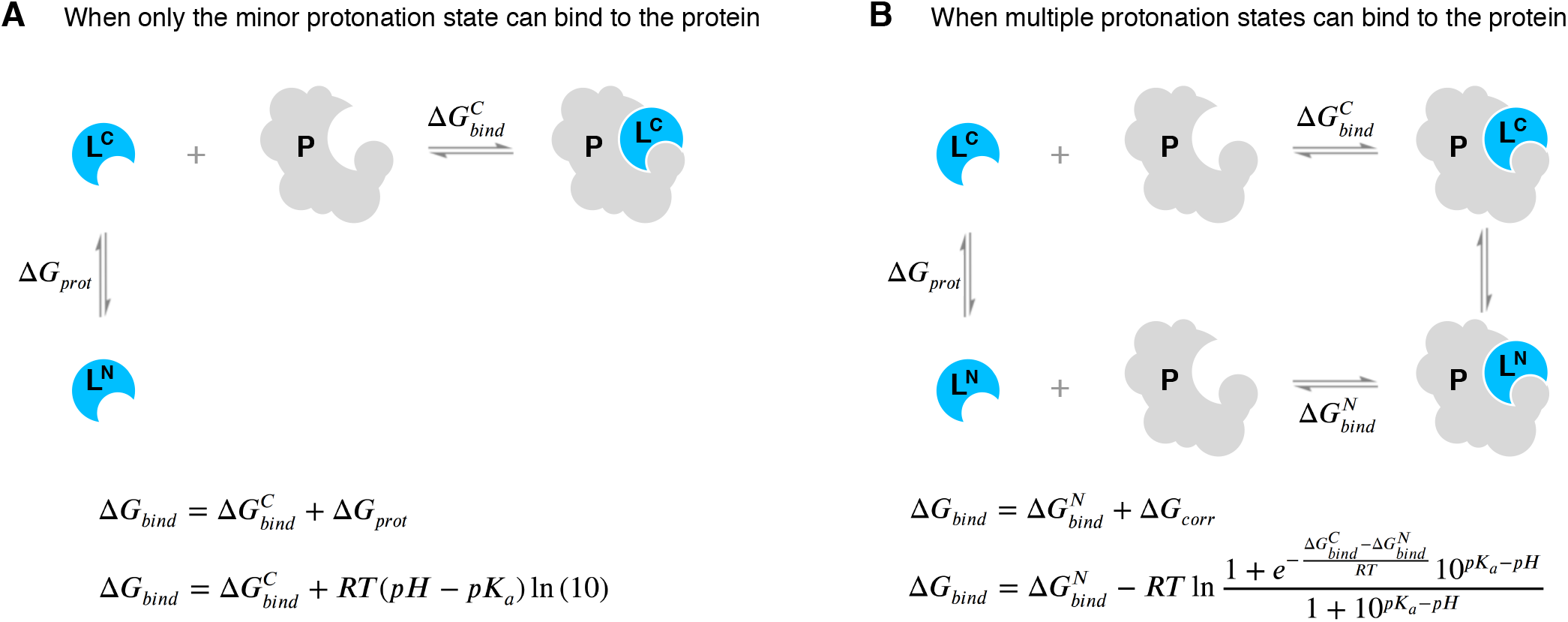
Aqueous ligand p*K*_a_ can influence overall protein-ligand binding affinity. **A** When only the minor aqueous protonation state contributes to protein-ligand complex formation, the overall binding free energy (Δ*G_bind_*) needs to be calculated as the sum of binding affinity of the minor state and the protonation penalty of that state. **B** When multiple charge states contribute to complex formation, the overall free energy of binding includes a multiple protonation states correction (MPSC) term (Δ*G_corr_*). MPSC is a function of pH, aqueous p*K*_a_ of the ligand, and the difference between the binding free energy of charged and neutral species 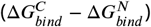.

In addition to the presence of multiple protonation states in the aqueous environment, multiple charge states can contribute to complex formation (Fig. 11B). Then, the overall free energy of binding needs to include a Multiple Protonation States Correction (MPSC) term (Δ*G_corr_*) [4]. MPSC is a function of pH, aqueous p*K*_a_ of the ligand, and the difference between the binding free energy of charged and neutral species 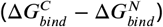 as shown in Fig. 11B.

Using the equations in Fig. 11B, we can model the true MPSC (Δ*G_corr_*)with respect to the difference between pH and the p*K*_a_ of the ligand to see when this value has a significant impact on the overall binding free energy. In Fig. 12, the true MPSC that must be added to 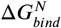 is shown for ligands with varying binding affinity difference between protonation states 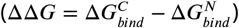. Fig. 12A shows the case of a monoprotic base in which the charged state has a lower affinity than the neutral state. Solid lines depict the accurate correction value. In cases where the p*K*_a_ is lower than the pH, the correction factor disappears as the ligand fully populates the neutral state 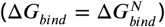. As the pH dips below the p*K*_a_, the charged state is increasingly populated and Δ*G_corr_* increases to approach ΔΔ*G*.

**Figure 12.**
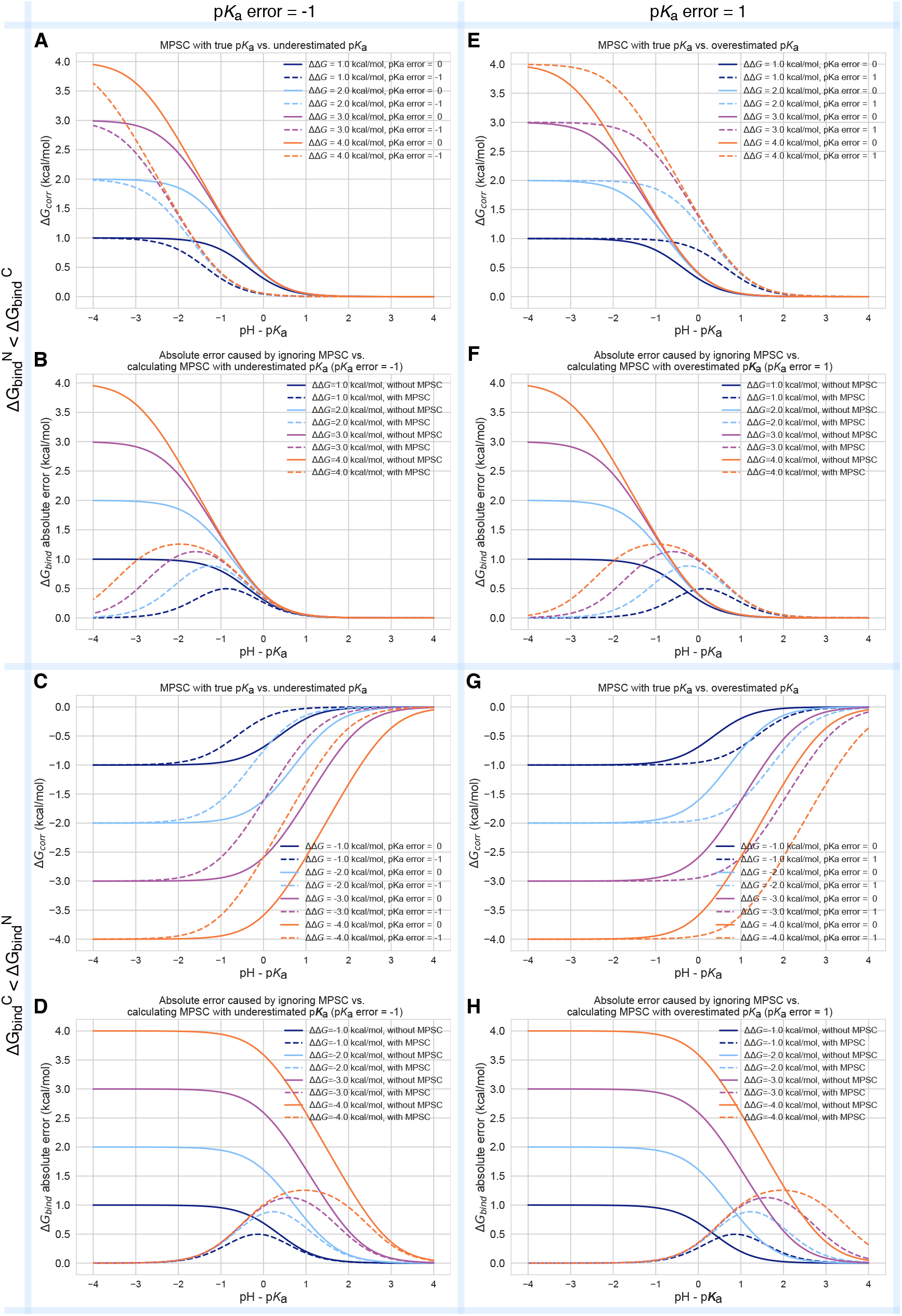
Inaccuracy of p*K*_a_ prediction (± 1 unit) affects the the accuracy of MPSC and overall protein-ligand binding free energy calculations to varying degrees based on aqueous p*K*_a_ and relative binding affinity of individual protonation states 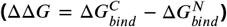. All calculations are made for 25°C, and a ligand with a single basic titratable group. **A, C, E, and G** show MPSC (Δ*G_corr_*) calculated with true vs. inaccurate p*K*_a_. **B, D, F, and H** show the comparison of the absolute error to Δ*G_bind_* caused by ignoring the MPSC completely (solid lines) vs. calculating MPSC based in inaccurate p*K*_a_ value (dashed lines). These plots provide guidance on when it is beneficial to include MPSC correction based on p*K*_a_ error, pH - p*K*_a_, and ΔΔ*G*.

It is interesting to note the pH-p*K*_a_ range over which Δ*G_corr_* changes significantly. It is often assumed that, for a basic ligand, if the p*K*_a_ of a ligand is more than 2 units higher than the pH, only 1% of the population is in the neutral state according to Henderson-Hasselbalch equation, and it is safe to approximate the overall binding affinity with 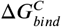. Based on the magnitude of the relative free energy difference between ligand protonation states, this assumption is not always correct. As seen in Fig. 12A, the responsive region of Δ*G_corr_* can span 3 pH units for a system with ΔΔ*G* = 1*kcal*/*mol*, or 5 pH units for a system with ΔΔ*G* = 4*kcal*/*mol*. This highlights that the range of p*K*_a_ values that impact binding affinity predictions is wider than 2 pH units. Molecules with p*K*_a_ values several units away from the physiological pH can still impact the overall binding affinity significantly due to the MPSC.

Despite the need to capture the contributions of multiple protonation states by including the MPSC in binding affinity calculations, inaccurate p*K*_a_ predictions can lead to errors in Δ*G_corr_* and overall free energy of binding prediction. In Fig. 12A dashed lines show predicted Δ*G_corr_* based on p*K*_a_ error of-1 units. We have chosen a p*K*_a_ error of 1 unit as this is the average inaccuracy expected from the p*K*_a_ prediction methods based on the SAMPL6 Challenge. Underestimation of the p*K*_a_ causes the Δ*G_corr_* to be underestimated as well and will result in overestimated affinities (i.e., too negative binding free energy) for a varying range of pH - p*K*_a_ values depending on the binding affinity difference between protonation states(ΔΔ*G*). In Fig. 12B dashed lines show how the magnitude of the absolute error caused by calculating Δ*G_corr_* with an inaccurate p*K*_a_ varies with respect to pH. Different colored lines show simulated results with varying binding free energy differences between protonation states. For a system whose charged state has higher binding free energy than the neutral state (ΔΔ*G* = 2 kcal/mol), the absolute error caused by underestimated p*K*_a_ by 1 unit can be up to 0.9 kcal/mol. For a system whose charged state has an even lower affinity (more positive binding free energy) than the neutral state (ΔΔ***G*** = 4 kcal/mol), the absolute error caused by underestimated p*K*_a_ by 1 unit can be up to 1.2 kcal/mol. The magnitude of errors contributing to overall binding affinity is too large to be neglected. Improving the accuracy of small molecule p*K*_a_ prediction methods can help to minimize the error in predicted MPSC.

With the current level of p*K*_a_ prediction accuracy as observed in SAMPL6 Challenge, is it advantageous to include the MPSC in affinity predictions that may include errors caused by p*K*_a_ predictions? We provide a comparison of the two choices to answer this question: (1) Neglecting the MPSC completely and assuming overall binding affinity is captured by 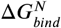, (2) including MPSC with a potential error in overall affinity calculation. The magnitude of error caused by Choice 1 (ignoring MPSC) is depicted as a solid line in Fig. 12B and the magnitude of error caused by MPSC computed with inaccurate p*K*_a_ is depicted as dashed lines. What is the best strategy? Error due to choice 1 is always larger than error due to choice 2 for all pH-p*K*_a_ values. In this scenario, including the MPSC improves overall binding affinity prediction accuracy. The error caused by the inaccurate p*K*_a_ is smaller than the error caused by neglecting the MPSC.

We can also ask whether or not an MPSC calculated based on an inaccurate p*K*_a_ should be included in binding affinity predictions in different circumstances, such as underestimated or overestimated p*K*_a_ values and charged states with higher or lower affinities than the neutral states. We tried to capture these circumstances in four quadrants of Fig. 12. In the case of overestimated p*K*_a_ values (Fig. 12E-H), it can be seen that for most of the pH-p*K*_a_ range, it is more advantageous to include the predicted MPSC in affinity calculations, except a smaller window where the opposite choice would be more advantageous. For instance, for the system with ΔΔ*G* = 2 kcal/mol and overestimated p*K*_a_ (Fig. 12E) for the pH-p*K*_a_ region between −0.5 and 2, including the predicted Δ*G_corr_* introduces more error than ignoring the MPSC.

In practice, we normally do not know the exact magnitude or the direction of the error of our predicted p*K*_a_. Therefore, using simulated MPSC error plots to decide when to include MPSC in binding affinity predictions is not possible. However, based on the analysis of a case with 1 unit of p*K*_a_ error, including the MPSC correction would be more often than not helpful in improving binding affinity predictions. The detrimental effect of p*K*_a_ inaccuracy is still significant. Hopefully, future improvements in p*K*_a_ prediction methodswill improve the accuracy of the MPSC and binding affinity predictions of ligands which have multiple protonation states that contribute to aqueous or complex populations. Being able to predict p*K*_a_ values with 0.5 units accuracy, for example, would significantly aid binding affinity models in computing more accurate MPSC terms.

The whole analysis presented in this section assumes that at least the dominant protonation state of the ligand is correctly included in the modeling of the protein-ligand complex. We have not discussed the case of omitting this dominant state from the free energy calculations entirely when it is erroneously predicted to be a minor state in solution. Such a mistake could be the most problematic, and the errors in estimated binding free energy could be very large.

### 3.4 Take-away lessons from SAMPL6 p*K*_a_ Challenge

The SAMPL6 p*K*_a_ Challenge showed that, in general, p*K*_a_ prediction accuracy of computational methods is lower than expected for drug-like molecules. Our expectation prior to the blind challenge was that well-developed methods would achieve prediction errors as low was 0.5 p*K*_a_ units, and make reliable predictions of dominant charge and tautomer states in solution. There are many factors that complicate predicting p*K*_a_ values of drug-like molecules: multiple titratable sites, including tautomerization, frequent presence of heterocycles, and extended conjugation patterns, as well as high numbers of rotatable bonds and the possibility of intramolecular hydrogen bonds. Macroscopic p*K*_a_ predictions have not yet reached experimental accuracy (where the inter-method variability of macroscopic p*K*_a_ measurements is around 0.5 p*K*_a_ units [23]). There was not a single method in the SAMPL6 Challenge that achieved RMSE around 0.5 or lower for macroscopic p*K*_a_ predictions for the 24 molecule set of kinase inhibitor fragment-like molecules. Smaller RMSEs were observed in the microscopic p*K*_a_ evaluation section of this study for some methods; however, the 8 molecule set used for that analysis poses a very limited dataset to reach conclusions about general expectations for drug-like molecules.

As the majority of experimental data was in the form of macroscopic p*K*_a_ values, we had to adopt a numerical matching algorithm (Hungarian matching) to pair predicted and experimental values to calculate performance statistics of macroscopic p*K*_a_ predictions. Accuracy, correlation, and extra/missing p*K*_a_ prediction counts were the main metrics for macroscopic p*K*_a_ evaluations. An RMSE range of 0.7 to 3.2 p*K*_a_ units was observed for all methods. Only five methods achieved RMSE between 0.7-1 p*K*_a_ units, while an RMSE between 1.5-3 log units was observed for the majority of methods. All four methods of the LFER category and three out of 5 QSPR/ML methods achieved RMSE less than 1.5 p*K*_a_ units. All the QM methods that achieved this level of performance included linear empirical corrections to rescale and unbias their p*K*_a_ predictions.

Based on the consideration of multiple error metrics, we compiled a shortlist of consistently-well performing methods for macroscopic p*K*_a_ evaluations. Two methods from QM+LEC methods, one QSPR/ML, two empirical methods achieved consistent performance according to many metrics. The common features of the two empirical methods were their large training sets (16000-17000 compounds) and commercial nature.

There were four submissions of QM-based methods that utilized the COSMO-RS implicit solvation model. While three of these achieved the lowest RMSE among QM-based methods (*xvxzd, yqkga*, and *8xt50*) [46], one of them showed the highest RMSE (*0hxtm* (COSMOtherm_FINE17)). The comparison of these methods indicates that capturing the conformational ensemble of microstates, using high-level QM calculations, and including RRHO corrections contribute to better macroscopic p*K*_a_ predictions. Linear empirical corrections applied QM calculations improved results, especially when the linear correction is calibrated for an experimental dataset using the same level of theory as the deprotonation free energy predictions (as in *xvxzd*). This challenge also points to the advantage of the COSMO-RS solvation approach compared to other implicit solvent models.

Molecules that posed greater difficulty for p*K*_a_ predictions were determined by comparing the macroscopic p*K*_a_ prediction accuracy of each molecule averaged over all methods submitted to the challenge. p*K*_a_ prediction errors were higher for compounds with sulfur-containing heterocycles, iodo, and bromo groups. This trend was also conserved when only QM-based methods were analyzed. The SAMPL6 p*K*_a_ dataset consisted of only 24 small molecules which limited our ability to statistically confirm this conclusion, however, we believe it is worth reporting molecular features that coincided with larger errors even if we can not evaluate the reason for these failures.

Utilizing a numerical matching algorithm to pair experimental and predicted macroscopic p*K*_a_ values was a necessity, however, this approach did not capture all aspects of prediction errors. Computing the number of missing or extra p*K*_a_ predictions remaining after Hungarian matching provided a window for observing macroscopic p*K*_a_ prediction errors such as the number of macroscopic transitions or ionization states expected in a pH interval. In p*K*_a_ evaluation studies, it is typical to just focus on p*K*_a_ value errors evaluated after matching and to ignore p*K*_a_ prediction errors that the matching protocol can not capture [49–53]. Frequently ignored prediction errors include predicting missing or extra p*K*_a_s and failing to predict the correct charge states. The SAMPL6 p*K*_a_ Challenge results showed sporadic presence of missing p*K*_a_ predictions and very frequent tendency to make extra p*K*_a_ predictions. Both indicate failures to capture the correct ionization states. The traditional way of evaluating p*K*_a_s that only focuses on the p*K*_a_ value error after some sort of numerical match between predictions and experimental values may have motivated these types of errors as there would be no penalty for missing a macroscopic deprotonation and predicting an extra one. This problem does not seem to be specific to any method category.

We used the eight molecule subset of SAMPL6 compounds with NMR-based dominant microstate sequence information to demonstrate the advantage of evaluating p*K*_a_ prediction on the level of microstates. Comparison of statistics computed for the 8 molecule dataset by Hungarian matching and microstate-based matching showed how Hungarian matching, despite being the best choice when only numerical matching is possible, can still mask errors in p*K*_a_ predictions. Errors computed by microstate-based matching were larger compared to numerical matching algorithms in terms of RMSE. Microscopic p*K*_a_ analysis with numerical matching algorithms may mask errors due to the higher number of guesses made. Numerical matching based on p*K*_a_ values also ignores information regarding the relative population of states. Therefore, it can lead to p*K*_a_s defined between very low energy microstate pairs to be matched to the experimentally observable p*K*_a_ between microstatesof higher populations. Of course, the predicted p*K*_a_ value could be correct however the predicted microstates would be wrong. Such mistakes caused by Hungarian matching were observed frequently in SAMPL6 results, and therefore we decided microstate-based matching of p*K*_a_values provides a more realistic picture of method performance.

Some QM and LFER methods made mistakes in predicting the dominant tautomers of the ionization states. Dominant tautomer prediction seemed to be particularly difficult for charged tautomers compared with neutral tautomers. The easiest way to extract the dominant microstate sequence from predictions was to calculate the relative free energy of microstates at any reference pH, determining the lowest free energy state in each ionization state. Errors in dominant microstate predictions were very rare for neutral tautomers, but more frequent in cationic tautomers with +1 charge of the 8 molecule set. SM14 was the molecule with the lowest dominant microstate prediction accuracy, while dominant microstates predictions for SM15 were perfect for all molecules. SM14 and SM15 both possess two experimental p*K*_a_s and a benzimidazole scaffold. The difference between them is the distance between the experimental p*K*_a_ values, which is smaller for SM14. These results make sense from the perspective of relative free energies of microstates. Closer p*K*_a_ values mean that the free energy difference between different microstates is smaller for SM14, and therefore any error in predicting the relative free energy of tautomers is more likely to cause reordering of relative populations of microstates and impact the accuracy of dominant microstate predictions. It would have been extremely informative to evaluate the tautomeric ratios and relative free energy predictions of microstates, however, the experimental data needed for this approach was not available. Tautomeric ratios could not be measured by the experimental methods available to us. Resolving tautomeric ratios would require extensive NMR measurements, but these measurements can suffer from lower accuracy especially when the free energy difference between tautomers is large.

The overall assessment of the SAMPL6 p*K*_a_ Challenge captured non-stellar performance for microscopic and macroscopic p*K*_a_ predictions which can be detrimental to the accuracy of protein-ligand affinity predictions and other pH-dependent physicochemical property predictions such as distribution coefficients, membrane permeability, and solubility. Protein-ligand binding affinity predictions utilize p*K*_a_ predictions in two ways: determination of the relevant aqueous microstates and quantification of the free energy penalty to reach these states. More accurate microscopic p*K*_a_ predictions are needed to be able to accurately incorporate multiple protonation state corrections (MPSC) into overall binding affinity calculations.

We simulated the effect of overestimating or underestimating p*K*_a_ of a ligand by one unit on overall binding affinity prediction for a ligand where both cation and neutral states contribute to binding affinity. A p*K*_a_ prediction error of this magnitude (assuming dominant tautomers were predicted correctly) could cause up to 0.9 and 1.2 kcal/mol error in overall binding affinity when the binding affinity of protonation states are 2 or 4 kcal/mol different, respectively. For the case of 4 kcal/mol binding affinity difference between protonation states, the pH-p*K*_a_ range that the error would be larger than 0.5 kcal/mol surprisingly spans around 3.5 pH units. The worse case, of course, is where there is a significant difference in binding free energy between the two protonation states, but we include the wrong one in our free energy calcuation. We demonstrated that the range of pH-p*K*_a_ value that the MPSC needs to be incorporated in binding affinity predictions can be wider than the widely assumed range of 2 pH units, based on the affinity difference between protonation states. At the level of 1 unit p*K*_a_ error, incorporating the MPSC would improve binding affinity predictions more often than not. If the microscopic p*K*_a_ could be predicted with 0.5 p*K*_a_ units of accuracy, MPSC calculations would be much more reliable.

There are multiple factors to consider when deciding which p*K*_a_ prediction method to utilize. These factors include the accuracy of microscopic and macroscopic p*K*_a_ values, the accuracy of the number and the identity of ionization states predicted within the experimental pH interval, the accuracy of microstates predicted within the experimental pH interval, the accuracy of tautomeric ratio (i.e., relative free energy between microstates), how costly is the calculation in terms of time and resources, and whether one has access to software licenses that might be required.

All of the top-performing empirical methods were developed as commercial software that requires a license to run, and there were not any open-source alternatives for empirical p*K*_a_ predictions. Since the completion of the blind challenge, two publications reported open-source machine learning-based p*K*_a_ prediction methods, however, one can only predict the most acidic or most basic macroscopic p*K*_a_ values of a molecule [54] and the second one is only trained for predicting p*K*_a_ values of monoprotic molecules [55]. Recently, a p*K*_a_ prediction methodology was published that describes a mixed approach of semi-empirical QM calculations and machine learning that can predict macroscopic p*K*_a_s of both mono- and polyprotic species [56]. The authors reported RMSE of 0.85 for the retrospective analysis performed on the SAMPL6 dataset.

### 3.5 Suggestions for future blind challenge design and evaluation of p*K*_a_ predictions

This analysis helped us understand the current state of the field and led to many lessons informing future SAMPL challenges. We believe the greatest benefit can be achieved if further iterations of small molecule p*K*_a_ prediction challenges can be organized, creating motivation for improving protonation state prediction methods for drug-like molecules. In future challenges, it is desirable to increase chemical diversity to cover more common scaffolds [57] and functional groups [58] seen in drug-like molecules, gradually increasing the complexity of molecules.

#### Microscopic p*K*_a_ measurements are needed for careful benchmarking of p*K*_a_ predictions for multiprotic molecules

Future challenges should promote stringent evaluation for p*K*_a_ prediction methods from the perspective of microscopic p*K*_a_ and microstate predictions. It is necessary to assess the capability of p*K*_a_ prediction methods to capture the free energy profile of microstates of multiprotic molecules. This is critical because p*K*_a_ predictions are often utilized to determine relevant protonation states and tautomers of small molecules that must be captured in other physical modeling approaches, such as protein-ligand binding affinity or distribution coefficient predictions. Different tautomers can have different binding affinities and partition coefficients.

In this paper, we demonstrated how experimental microstate information can guide the analysis further than the typical p*K*_a_ evaluation approach that has been used so far. The traditional p*K*_a_ evaluation approach focuses solely on the numerical error of the p*K*_a_ values and neglects the difference between macroscopic and microscopic p*K*_a_ definitions. This is mainly caused by the lack of p*K*_a_ datasets with microscopic detail. To improve p*K*_a_ and protonation state predictions for multiprotic molecules, it is necessary to embrace the difference between macroscopic and microscopic p*K*_a_ definitions and select strategies for experimental data collection and prediction evaluation accordingly. In the SAMPL6 Challenge, the analysis was limited by the availability of experimental microscopic data as well. As is usually the case, macroscopic p*K*_a_ values were abundant (24 molecules) and limited data on microscopic states was available (8 molecules), although the latter opened new avenues for evaluation. For future blind challenges for multiprotic compounds, striving to collect experimental datasets with microscopic p*K*_a_s would be very beneficial, despite the high cost of these measurements. Benchmark datasets of microscopic p*K*_a_ values with assigned microstates are currently missing because experimental determination of these are much more expensive and time-consuming than macroscopic p*K*_a_ measurements. This limits the ability to improve p*K*_a_ and tautomer prediction methods for multiprotic molecules. If the collection of experimental microscopic p*K*_a_s is not possible due to time and resource costs of such NMR experiments, at least supplementing the more automated macroscopic p*K*_a_ measurements with NMR-based determination of the dominant microstate sequence or tautomeric ratios of each ionization state can create very useful benchmark datasets. This supplementary information can allow microstate-based assignment of experimental to predicted p*K*_a_ values and a more realistic assessment of method performance.

#### Evaluation strategy for p*K*_a_ predictions must be determined based on the nature of experimental p*K*_a_ measurements available

If the only available experimental data is in the form of macroscopic p*K*_a_ values, the best way to evaluate computational predictions is by calculating predicted macroscopic p*K*_a_ from microscopic p*K*_a_ predictions. With the conversion of microscopic p*K*_a_ to macroscopic p*K*_a_s, all structural information about the titration site is lost, and the only remaining information is the total charge of macroscopic ionization states. Unfortunately, most macroscopic p*K*_a_ measurements—including potentiometric and spectrophotometric methods—do not capture the absolute charge of the macrostates. The spectrophotometric method does not measure charge at all. The potentiometric method can only capture the relative charge changes between macrostates. Only pH-dependent solubility-based p*K*_a_ estimations can differentiate neutral and charged states from one another. It is, therefore, very common to have experimental datasets of macroscopic p*K*_a_ without any charge or protonation position information regarding the macrostates. This causes an issue of assigning predicted and experimental p*K*_a_ values before any error statistics can be calculated.

As delineated by Fraczkiewicz [23], the fairest and most reasonable solution for the p*K*_a_ matching problem involves an assignment algorithm that preserves the order of predicted and experimental microstates and uses the principle of smallest differences to pair values. We recommend Hungarian matching with a squared-error penalty function. The algorithm is available in SciPy package (scipy.optimize.linear_sum_assignment) [35]. In addition to the analysis of numerical error statistics following Hungarian matching, at the very least, the number of missing and extra p*K*_a_ predictions must be reported based on unmatched p*K*_a_ values. Missing or extra p*K*_a_ predictions point to a problem with capturing the right number of ionization states within the pH interval of the experimental measurements. We have demonstrated that for microscopic p*K*_a_ predictions, performance analysis based on Hungarian matching results in overly optimistic and misleading results—instead the employed microstate-based matching provided a more realistic assessment when microstate data is available.

#### Lessons from the first p*K*_a_ blind challenge will guide future decisions on challenge rules, prediction reporting formats, and challenge inputs

We solicited three different submission types in SAMPL6 to capture all the necessary information related to p*K*_a_ predictions. These were (1) macroscopic p*K*_a_ values, (2) microscopic p*K*_a_ values and microstate pair identities, and (3) fractional population of microstates with respect to pH. We realized later that collecting fractional populations of microstates was redundant since microscopic p*K*_a_ values and microstate pairs capture all the necessary information to construct fractional population vs. pH curves [26]. Only microscopic and macroscopic p*K*_a_ values were used for the challenge analysis presented in this paper.

While exploring ways to evaluate SAMPL6 p*K*_a_ Challenge results, we developed a better way to capture microscopic p*K*_a_ predictions, as presented in Gunner et al. [26]. This alternative reporting format consists of reporting the charge and relative free energy of microstates with respect to an arbitrary reference microstate and pH. This approach presents the most concise method of capturing all necessary information regarding microscopic p*K*_a_ predictions and allows calculation of predicted microscopic p*K*_a_s, microstate population with respect to pH, macroscopic p*K*_a_ values, macroscopic population with respect to pH, and tautomer ratios. Still, there may be methods developed to predict macroscopic p*K*_a_s directly instead of computing them from microstate predictions that justifies allowing a macroscopic p*K*_a_ reporting format. In future challenges, we recommend collecting p*K*_a_ predictions with two submission types: (1) macroscopic p*K*_a_ values together with the charges of the macrostates and (2) microstates, their total charge, and relative free energies with respect to a specified reference microstate and pH. This approach is being used in SAMPL7.

In SAMPL6, we provided an enumerated list of microstates and their assigned microstate IDs because we were worried about parsing submitted microstates in SMILES from different sources correctly. There were two disadvantages to this approach. First, this list of enumerated microstates was used as input by some participants which was not our intention. (Challenge instructions requested that predictions should not rely on these microstate lists and only use them for matching microstate IDs.) Second, the first iteration of enumerated microstates was not complete. We had to add new microstates and assign them microstate IDs for a couple of rounds until reaching a complete list. In future challenges, a better way of handling the problem of capturing predicted microstates would be asking participants to specify the predicted protonation states themselves and assigning identifiers after the challenge deadline to aid comparative analysis. This would prevent the partial unblinding of protonation states and allow the assessment of whether methods can predict all the relevant states independently, without relying on a provided list of microstates. Predicted states can be submitted as mol2 files that represent the microstate with explicit hydrogens. The organizers must only provide the microstate that was selected as the reference state for the relative microstate free energy calculations.

In the SAMPL6 p*K*_a_ Challenge, there was not a requirement that participants should report predictions for all compounds. Some participants reported predictions for only a subset of compounds, which may have led these methods to look more accurate than others due to missing predictions. In the future, it will be better to allow submissions of only complete sets for a better comparison of method performance.

A wide range of methods participated in the SAMPL6 p*K*_a_ Challenge—from very fast QSPR methods to QM methods with a high-level of theory and extensive exploration of conformational ensembles. In the future, it would be interesting to capture computing costs in terms of average compute hours per molecule. This can provide guidance to future users of p*K*_a_ prediction methods for selection of which method to use.

#### It is advantageous to field associated challenges with common set of molecules for different physicochemical properties

Future blind challenges can maximize learning opportunities by evaluating predictions of different physicochemical properties for the same molecules in consecutive challenges. In SAMPL6, we organized both p*K*_a_ and log *P* challenges. Unfortunately only a subset of compounds in the p*K*_a_ datasets were suitable for the potentiometric log *P* measurements [8]. Still, comparing prediction performance of common compounds in both challenges can lead to beneficial insights especially for physical modeling techniques if there are common aspects that are beneficial or detrimental to prediction performance. For example, in SAMPL6 p*K*_a_ and log *P* Challenges COSMO-RS and EC-RISM solvation models achieved good performance. Having access to a variety of physicochemical property measurements can also help the identification of error sources. For example, dominant microstates determined for p*K*_a_ challenge can provide information to check if correct tautomers are modeling in a log *P* or log *D* challenge. p*K*_a_ prediction is a requirement for log *D* prediction and experimental p*K*_a_ values can help diagnosing the source of errors in log *D* predictions better. The physical challenges in SAMPL7, for which the blind portion of the challenges have just concluded on October 8th, 2020, follow this principle and include both p*K*_a_, log *P*, and membrane permeability properties for a set of mono-protic compounds. We hope that future p*K*_a_ challenges can focus on multiprotic drug-like compounds with microscopic p*K*_a_ measurements for an in-depth analysis.

## 4 Conclusion

The first SAMPL6 p*K*_a_ Challenge focused on molecules resembling fragments of kinase inhibitors, and was intended to assess the performance of p*K*_a_ predictions for drug-like molecules. With wide participation, we had an opportunity to prospectively evaluate p*K*_a_ predictions spanning various empirical and QM based approaches. In addition to community participants, a small number of popular p*K*_a_ prediction methods that were missing from blind submissions were added as reference calculations after the challenge deadline.

Practical experimental limitations restricted the overall size and microscopic information available for the blind challenge dataset [8]. The experimental dataset consisted of spectrophotometric measurements of 24 molecules, some of which were multiprotic. For a subset of molecules there was also NMR data to inform the dominant microstate sequence, though microscopic p*K*_a_ measurements were not performed. We conducted a comparative analysis of methods represented in the blind challenge in terms of both macroscopic and microscopic p*K*_a_ prediction performance avoiding any assumptions about the interpretation of experimental p*K*_a_s.

Here, we used Hungarian matching to assign predicted and experimental values for the calculation of accuracy and correlation statistics, because the majority of experimental data was macroscopic p*K*_a_ values. In addition to evaluating error in predicted p*K*_a_ values, we also reported the macroscopic p*K*_a_ errors that were not captured by the match between experimental and predicted p*K*_a_ values. These were extra or missing p*K*_a_ predictions which are important indicators that predictions are failing to capture the correct ionization states.

We evaluated microscopic p*K*_a_ predictions utilizing the experimental dominant microstate sequence data of eight molecules. This experimental data allowed us to use microstate-based matching for evaluating the accuracy of microscopic p*K*_a_ values in a more realistic way. We have determined that QM and LFER predictions had lower accuracy in determining the dominant tautomer of the charged microstates than the neutral states. For both macroscopic and microscopic p*K*_a_ predictions we have determined methods that were consistently well-performing according to multiple statistical metrics. Focusing on the comparison of molecules instead of methods for macroscopic p*K*_a_ prediction accuracy indicated molecules with sulfur-containing heterocycles, iodo, and bromo groups suffered from lower p*K*_a_ prediction accuracy.

The overall performance of p*K*_a_ predictions as captured in this challenge is concerning for the application of p*K*_a_ prediction methods in computer-aided drug design. Many computational methods for predicting target affinities and physicochemical properties rely on p*K*_a_ predictions for determining relevant protonation states and the free energy penalty of such states. 1 unit of p*K*_a_error is an optimistic estimate of current macroscopic p*K*_a_ predictions for drug-like molecules based on SAMPL6 Challenge where errors in predicting the correct number of ionization states or determining the correct dominant microstate were also common to many methods. In the absence of other sources of errors, we showed that 1 unit over- or underestimation of the p*K*_a_ of a ligand can cause significant errors in the overall binding affinity calculation due to errors in multiple protonation state correction factor.

The SAMPL6 GitHub Repository contains all information regarding the challenge structure, experimental data, blind prediction submission sets, and evaluation of methods. The repository will be useful for future follow up analysis and the experimental measurements can continue to serve as a benchmark dataset for testing methods.

In this article, we aimed to demonstrate not only the comparative analysis of the p*K*_a_ prediction performance of contemporary methods for drug-like molecules, but also to propose a stringent p*K*_a_ prediction evaluation strategy that takes into account differences in microscopic and macroscopic p*K*_a_ definitions. We hope that this study will guide and motivate further improvement of p*K*_a_ prediction methods.

## Supporting information

supplementary-documents.tar.gz

## Abbreviations

SAMPL: Statistical Assessment of the Modeling of Proteins and Ligands
p*K*_a_: log_10_ of the acid dissociation equilibrium constant
log *P*: log_10_ of the organic solvent-water partition coefficient (***K***_*ow*_) of neutral species
log *D*: log_10_ of organic solvent-water distribution coefficient (***D***_*ow*_)
SEM: Standard error of the mean
RMSE: Root mean squared error
MAE: Mean absolute error
*τ*: Kendall’s rank correlation coefficient (Tau)
R^2^: Coefficient of determination (R-Squared)
MPSC: Multiple protonation states correction for binding free energy
DL: Database Lookup
LFER: Linear Free Energy Relationship
QSPR: Quantitative Structure-Property Relationship
ML: Machine Learning
QM: Quantum Mechanics
LEC: Linear Empirical Correction

## 5 Code and data availability

- SAMPL6 p*K*_a_ challenge instructions, submissions, experimental data and analysis is available at SAMPL6 GitHub Repository: https://github.com/samplchallenges/SAMPL6

## 6 Overview of supplementary information

### Contents of the Supplementary Information

- TABLE S1: SMILES and InChI identifiers of SAMPL6 p*K*_a_ Challenge molecules.
- TABLE S2: Evaluation statistics calculated for all macroscopic p*K*_a_ prediction submissions based on Hungarian match for 24 molecules.
- TABLE S3: Evaluation statistics calculated for all microscopic p*K*_a_ prediction submissions based on Hungarian match for 8 molecules with NMR data.
- TABLE S4: Evaluation statistics calculated for all microscopic p*K*_a_ prediction submissions based on microstate match for 8 molecules with NMR data.
- FIGURE S1: Dominant microstates of 8 molecules were determined based on NMR measurements.
- FIGURE S2: MAE of macroscopic p*K*_a_ predictions of each molecule did not show any significant correlation with any molecular descriptor.
- FIGURE S3: The value of macroscopic p*K*_a_ was not a factor affecting prediction error seen in SAMPL6 Challenge according to the analysis with Hungarian matching.
- FIGURE S4: There was low agreement between experimental dominant microstate pairs and the predicted microstate pairs selected by Hungarian algorithm for microscopic p*K*_a_ predictions.

### Extra files included in *supplementary-documents.tar.gz*

- An archive copy of the p*K*_a_ Challenge directory of SAMPL6 GitHub Repository (*SAMPL6-repository-pKa-directory.zip*)
- Table S1 in CSV format (*SAMPL6-pKa-chemical-identifiers-table.csv*)
- Table S2 in CSV format (*macroscopic-pKa-statistics-24mol-hungarian-match.csv*)
- Table S3 in CSV format (*microscopic-pKa-statistics-8mol-hungarian-match-table.csv*)
- Table S4 in CSV format (*microscopic-pKa-statistics-8mol-microstate-match-table.csv*)
- Figure S1 in CSV format (*experimental-microstates-of-8mol-based-on-NMR.csv*)
- TheJupyter Notebook used for the enumeration of microstates (*enumerate-microstates-with-Epik-and-OpenEye-QUACPAC.ipynb*)
- A CSV table of SAMPL6 molecule IDs and OpenEye OEChem generated SMILES (*molecule_ID_and_SMILES.csv*)

## 7 Author Contributions

Conceptualization, MI, JDC; Methodology, MI, JDC, ASR; Software, MI, AR, ASR; Formal Analysis, MI, ASR; Investigation, MI; Resources, JDC, DLM; Data Curation, MI; Writing-Original Draft, MI; Writing - Review and Editing, MI, JDC, ASR, AR, DLM, MRG; Visualization, MI, AR; Supervision, JDC, DLM; Project Administration, MI; Funding Acquisition, JDC, DLM, MI.

## 8 Acknowledgments

We would like to acknowledge the infrastructure and website support of Mike Chiu that allowed a seamless collection of challenge submissions. Mike Chiu also provided assistance with constructing a submission validation script to ensure all submissions adhered to the machine-readable format. We are grateful to Kiril Lanevskij for suggesting the Hungarian algorithm for matching experimental and predicted p*K*_a_ values. We would like to thank Thomas Fox for providing MoKa reference calculations. We acknowledge Caitlin Bannan for guidance on defining a working microstate definition for the challenge and guidance for designing the challenge. We thank Brad Sherborne for his valuable insights at the conception of the p*K*_a_ challenge and connecting us with Timothy Rhodes and Dorothy Levorse who were able to provide resources and expertise for experimental measurements performed at MRL. We acknowledge Paul Czodrowski who provided feedback on multiple stages of this work: challenge construction, purchasable compound selection, and manuscript draft. MI,JDC, and DLM gratefully acknowledge support from NIH grant R01GM124270 supporting the SAMPL Blind Challenges. MI, ASR, AR, andJDC acknowledge support from the Sloan Kettering Institute. JDC acknowledges support from NIH grant P30CA008748 and NIH grant R01GM121505. DLM appreciates financial support from the National Institutes of Health (R01GM108889) and the National Science Foundation (CHE 1352608). MI acknowledges Doris J. Hutchinson Fellowship. MI, ASR, AR, andJDC are grateful to OpenEye Scientific for providing a free academic software license for use in this work. MI, ASR, AR, and JDC thank Janos Fejervari and ChemAxon team that gave us permission to include ChemAxon/Chemicalize p*K*_a_ predictions as a reference prediction in challenge analysis.

## 9 Disclaimers

The content is solely the responsibility of the authors and does not necessarily represent the official views of the National Institutes of Health.

## 10 Disclosures

JDC was a member of the Scientific Advisory Board for Schrödinger, LLC during part of this study, and is a current Scientific Advisory Board member for OpenEye Scientific and scientific advisor to Foresite Labs. DLM is a current member of the Scientific Advisory Board of OpenEye Scientific and an Open Science Fellow with Silicon Therapeutics.

The Chodera laboratory receives or has received funding from multiple sources, including the National Institutes of Health, the National Science Foundation, the Parker Institute for Cancer Immunotherapy, Relay Therapeutics, Entasis Therapeutics, Vir Biotechnology, Silicon Therapeutics, EMD Serono (Merck KGaA), AstraZeneca, Vir Biotechnology, XtalPi, the Molecular Sciences Software Institute, the Starr Cancer Consortium, the Open Force Field Consortium, Cycle for Survival, a Louis V. Gerstner Young Investigator Award, The Einstein Foundation, and the Sloan Kettering Institute. A complete list of funding can be found at http://choderalab.org/funding.

## 11 Supplementary Information

**Table S1.**
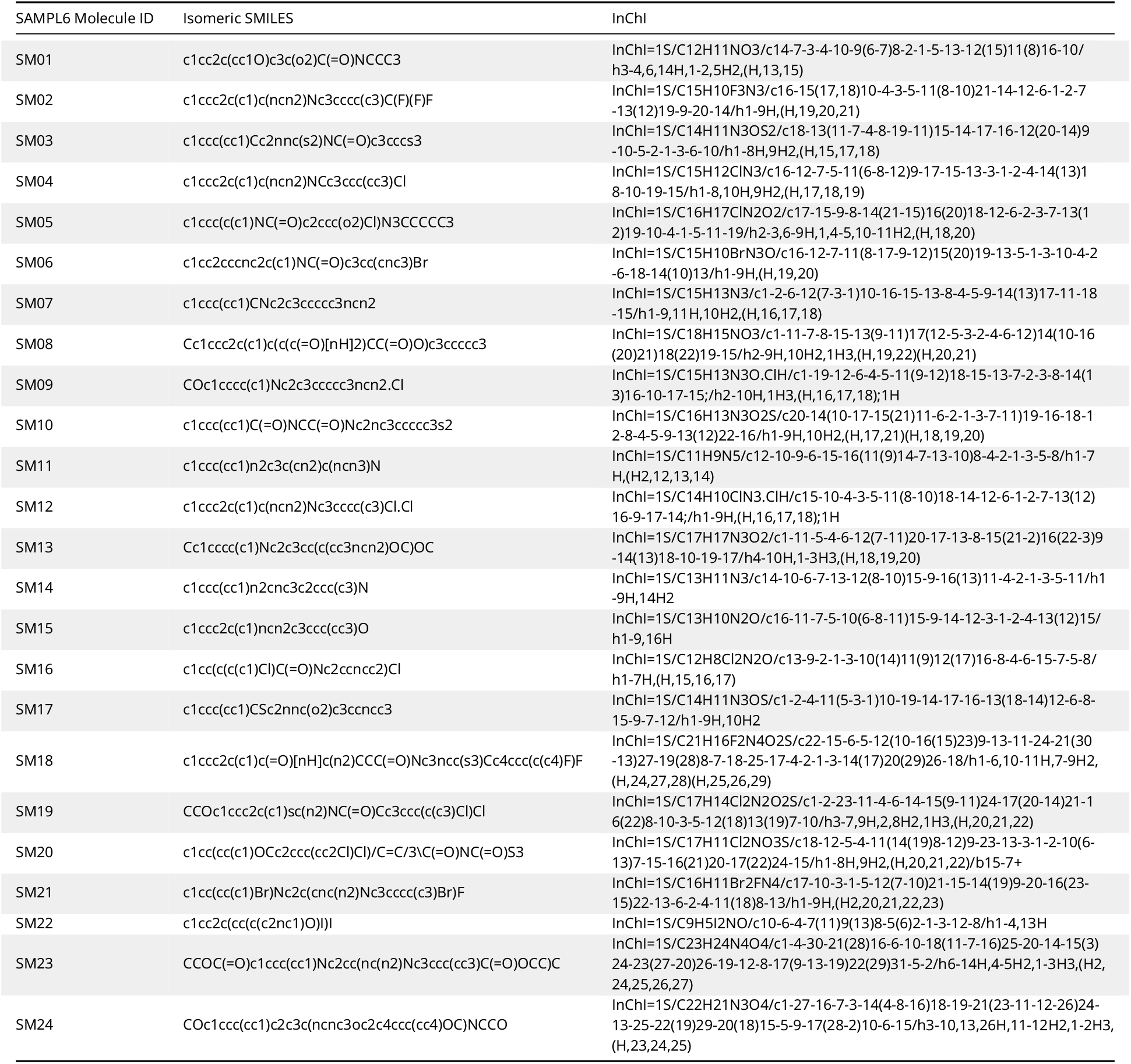
SMILES and InChI identifiers of SAMPL6 p*K*_a_ Challenge molecules. A CSV version of this table can be found in *SAMPL6-supplementary-documents.tar.gz*. SMILES were generated by OpenEye OEChem [32]

**Figure S1.**
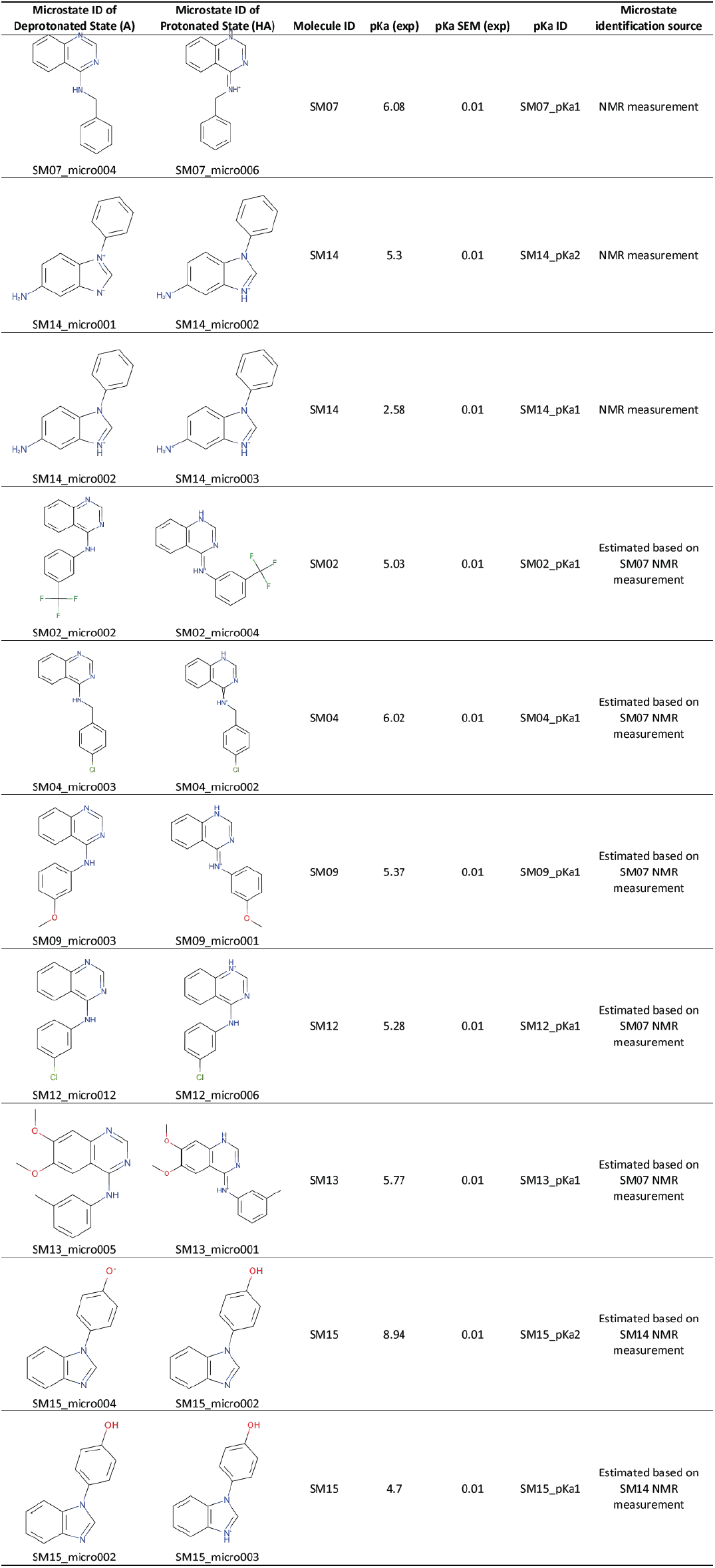
Dominant microstates of 8 molecules were determined based on NMR measurements. Dominant microstate sequence of 6 analogues were determined taking SM07 and SM14 as reference. Matched experimental p*K*_a_ values were determined by spectrophotometric p*K*_a_ measurements [8]. A CSV version of this table can be found in *SAMPL6-supplementary-documents.tar.gz*.

**Figure S2.**
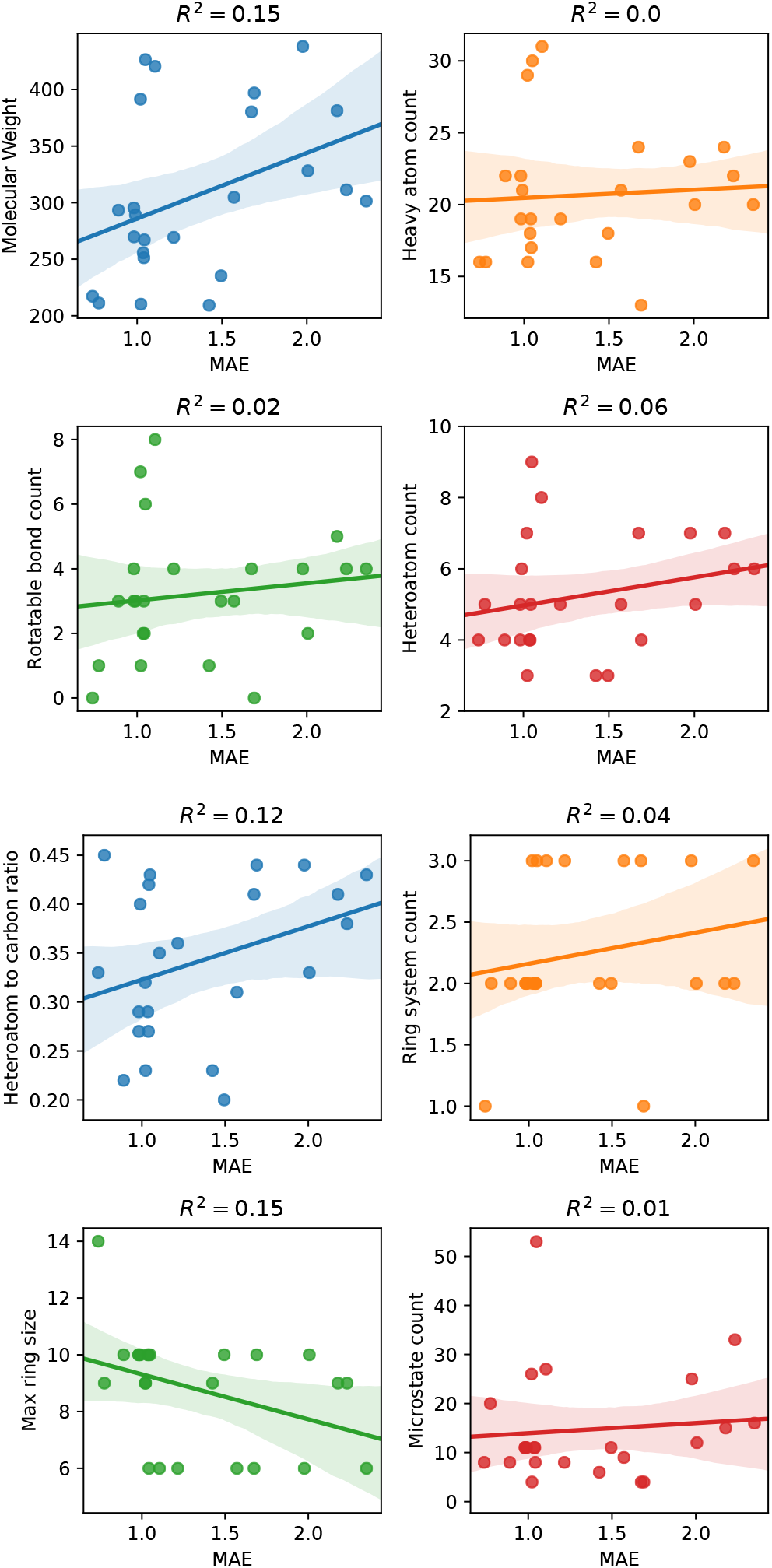
MAE of macroscopic p*K*_a_ predictions of each molecule did not show any significant correlation with any molecular descriptor. Plots show regression lines, 95% confidence intervals of the regression lines, and R_2_. The following molecular descriptors were calculated using OpenEye OEMolProp Toolkit [59]: molecular weight, non-terminal rotatable bond count, heteroatom to carbon ratio, maximum ring size, heavy atom count, heteroatom count, ring system count. Microstate count is based on the enumerated microstates for each compounds including additional microstates requested by participants.

**Figure S3.**
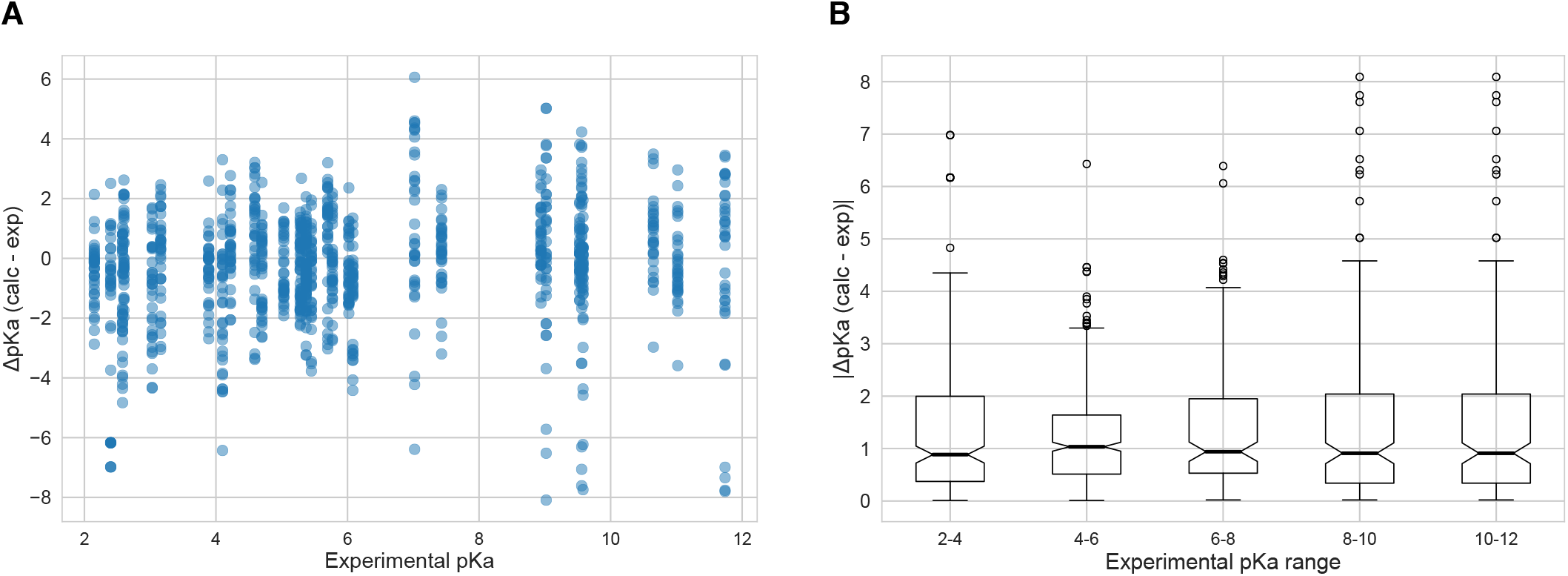
The value of macroscopic p*K*_a_s was not a factor affecting prediction error seen in SAMPL6 Challenge according to the analysis with Hungarian matching. There was not clear trend between p*K*_a_ prediction error and the true p*K*_a_ error. Very high and very low p*K*_a_ values have similar inaccuracy compared to p*K*_a_ values close to 7. **A** Scatter plot of macroscopic p*K*_a_ prediction error calculated with Hungarian matching vs. experimental p*K*_a_ value **B** Box plot of absolute error of macroscopic p*K*_a_ predictions binned into 2 p*K*_a_ unit intervals of experimental p*K*_a_.

**Figure S4.**
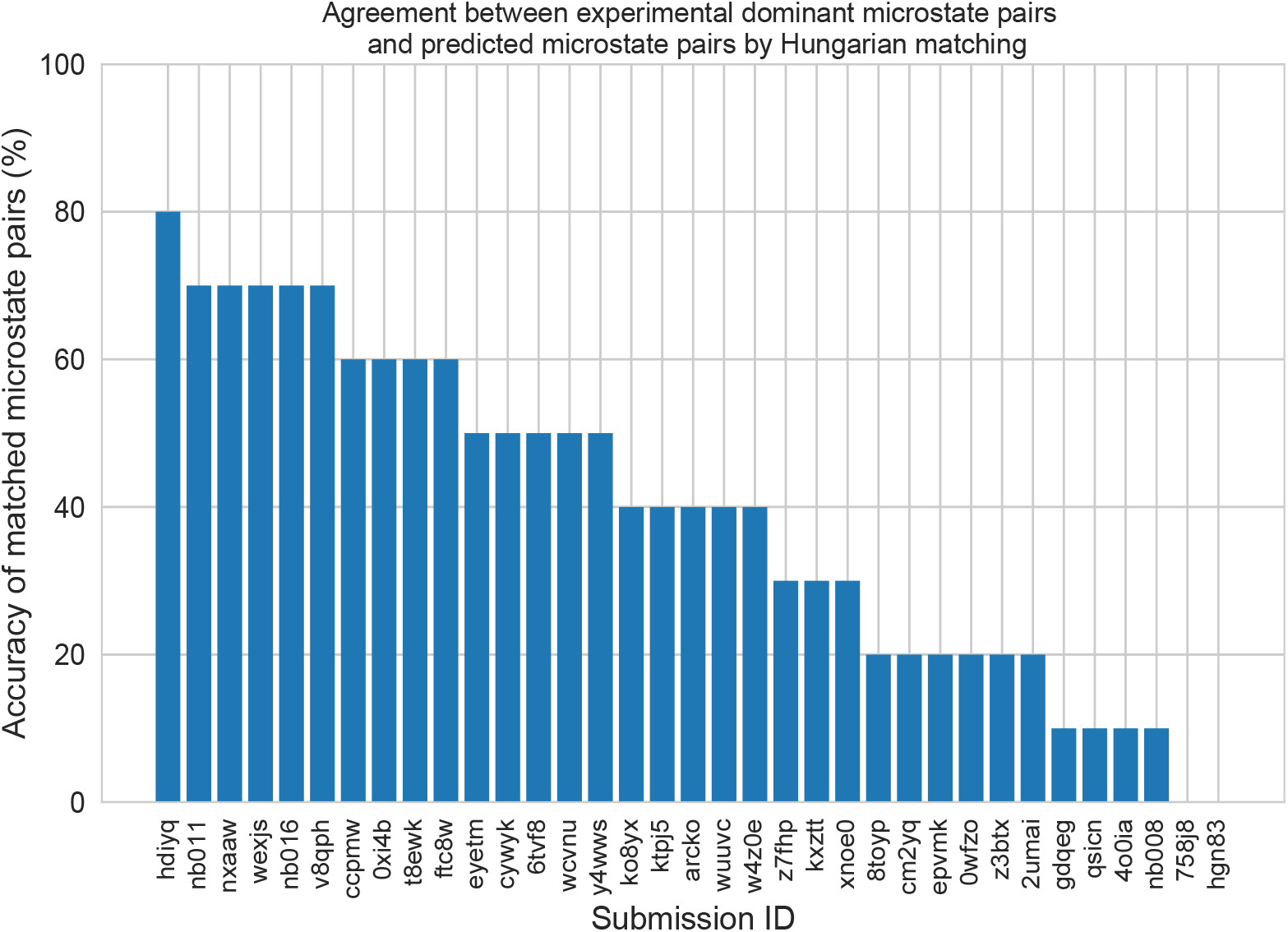
There was low agreement between experimental dominant microstate pairs and the predicted microstate pairs selected by Hungarian algorithm for microscopic p*K*_a_ predictions. This analysis could only be performed for 8 molecules with NMR data. Hungarian matching algorithm which matches predicted and experimental values considering only the closeness of the numerical value of p*K*_a_ and it often leads to predicted p*K*_a_ matches that described a different microstates pair than the experimentally observed dominant microstates.

**Table S2.**
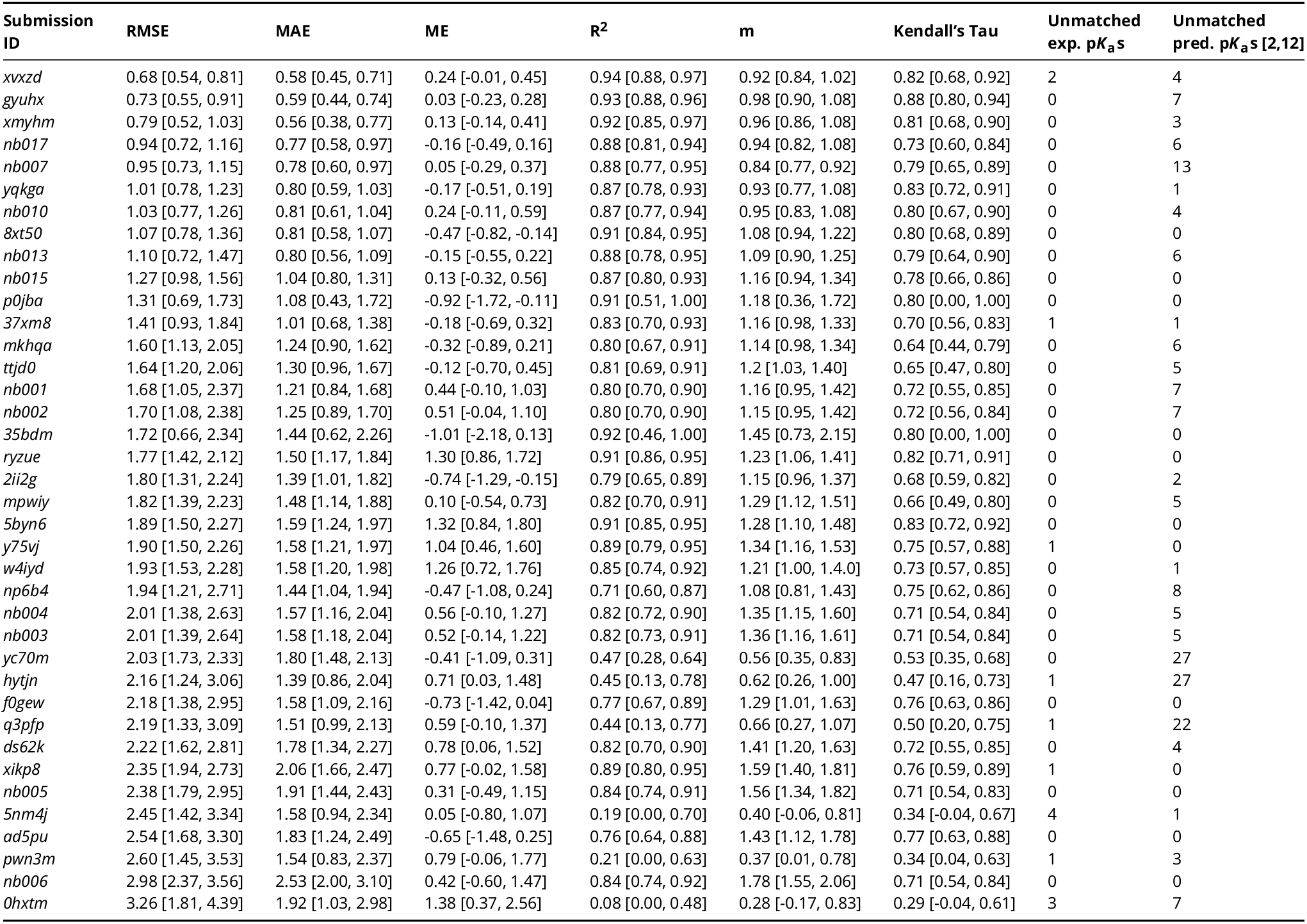
Evaluation statistics calculated for all macroscopic p*K*_a_ prediction submissions based on Hungarian match for 24 molecules. Methods are represented via their SAMPL6 submission IDs which can be cross-referenced with Table 1 for method details. There are eight error metrics reported: the root-mean-squared error (RMSE), mean absolute error (MAE), mean (signed) error (ME), coefficient of determination (R^2^), linear regression slope (m), Kendall’s Rank Correlation Coefficient (*τ*), unmatched experimental p*K*_a_s (number of missing p*K*_a_ predictions) and unmatched predicted p*K*_a_s (number of extra p*K*_a_ predictions between 2 and 12. This table is ranked by increasing RMSE. A CSV version of this table can be found in *SAMPL6-supplementary-documents.tar.gz*.

**Table S3.**
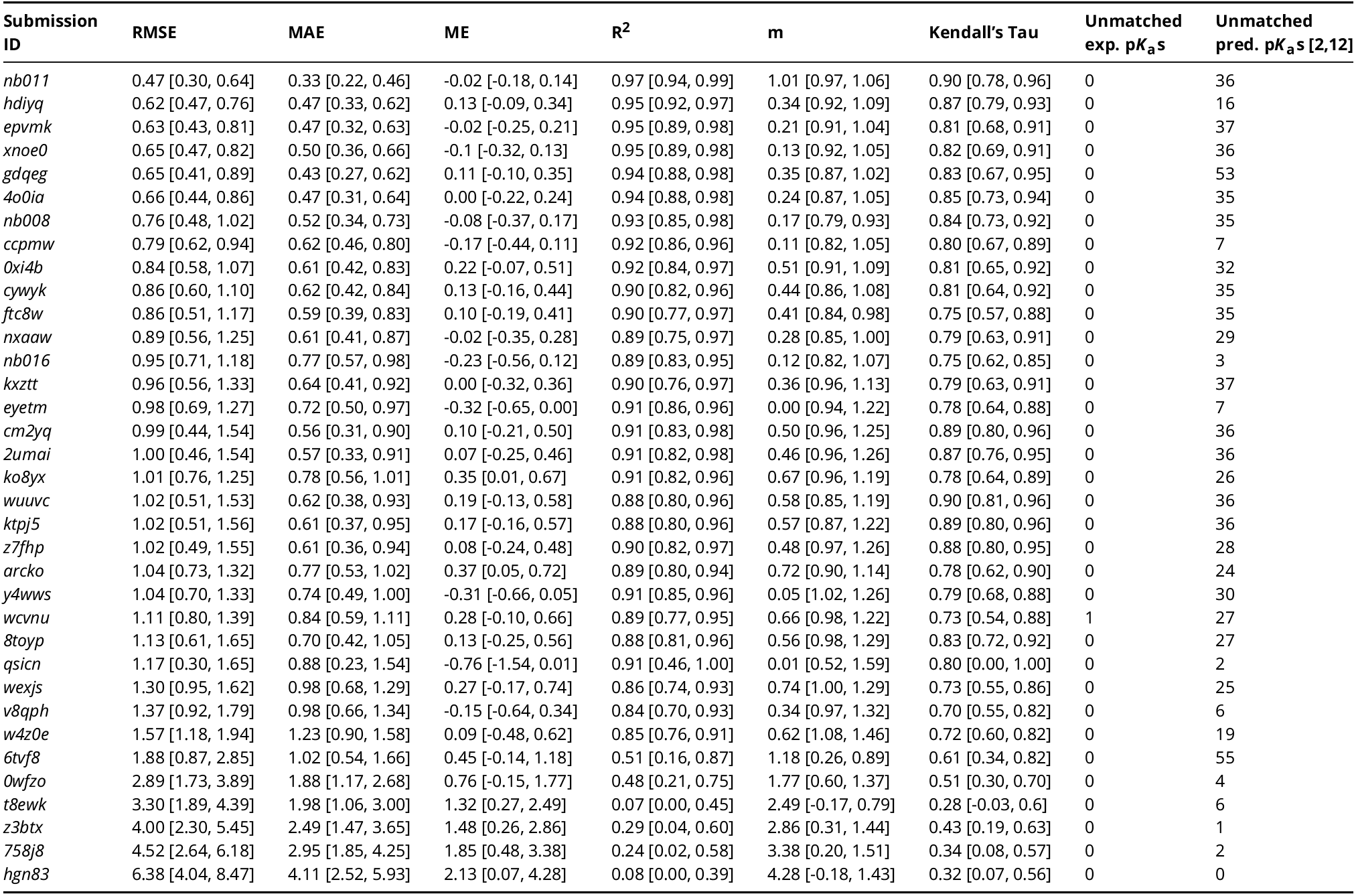
Evaluation statistics calculated for all microscopic p*K*_a_ prediction submissions based on Hungarian match for 8 molecules with NMR data. Methods are represented via their SAMPL6 submission IDs which can be cross-referenced with Table 1 for method details. There are eight error metrics reported: the root-mean-squared error (RMSE), mean absolute error (MAE), mean (signed) error (ME), coefficient of determination (R^2^), linear regression slope (m), Kendall’s Rank Correlation Coefficient (*τ*), unmatched experimental p*K*_a_s (number of missing p*K*_a_ predictions) and unmatched predicted p*K*_a_s (number of extra p*K*_a_ predictions between 2 and 12. This table is ranked by increasing RMSE. A CSV version of this table can be found in *SAMPL6-supplementary-documents.tar.gz*.

**Table S4.**
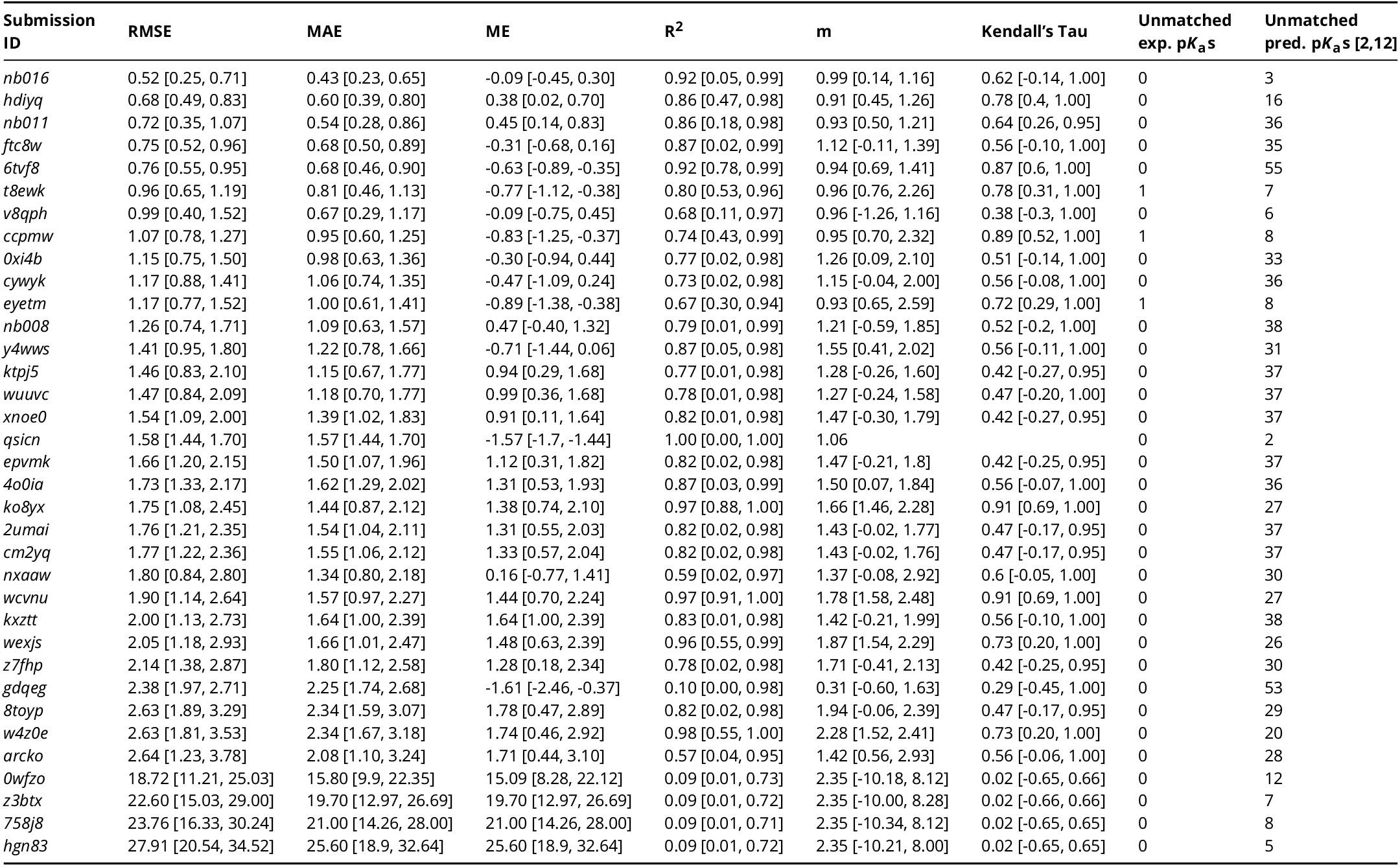
Evaluation statistics calculated for all microscopic p*K*_a_ prediction submissions based on microstate pair match for 8 molecules with NMR data. Methods are represented via their SAMPL6 submission IDs which can be cross-referenced with Table 1 for method details. There are eight error metrics reported: the root-mean-squared error (RMSE), mean absolute error (MAE), mean (signed) error (ME), coefficient of determination (R^2^), linear regression slope (m), Kendall’s Rank Correlation Coefficient (*τ*), unmatched experimental p*K*_a_s (number of missing p*K*_a_ predictions) and unmatched predicted p*K*_a_s (number of extra p*K*_a_ predictions between 2 and 12. This table is ranked by increasing RMSE. A CSV version of this table can be found in *SAMPL6-supplementary-documents.tar.gz*.

## Notes

https://github.com/samplchallenges/SAMPL6

