## supplementary-documents.tar.gz for "Overview of the SAMPL6 p*K*_a_ Challenge: Evaluating small molecule microscopic and macroscopic p*K*_a_ predictions": enumerate-microstates-with-Epik-and-OpenEye-QUACPAC.ipynb

```
# %% [markdown]
# Generating microstates for all molecules

Microstates are defined as tautomer and protomers of a molecule. We enumerated microstates with Epik and OpenEye Quakpac and took the union of results. Even though resonance structures correspond to different canonical isomeric SMILES they are not different microstates. Multiple resonance structures were removed. 

To detect multiple resonance structures we compared fixed hydrogen InChI strings. Resonance structures lead to same InChI string but tautomers and protomers have unique InChI strings. 

# %% [code]
import pandas as pd
import numpy as np
import os, sys
from openeye.oechem import *
from openeye.oequacpac import *

def prepare_inchi(mol):
    """Create InChI hashes with explicit, fixed hydrogen layer from an OEGraphMol."""
    opts = OEInChIOptions()
    # Do not add hydrogens.
    opts.SetHydrogens(False)
    # Return the fixed hydrogen layer
    opts.SetFixedHLayer(True)
    return OECreateInChI(mol, opts)

# SAMPL6 small molecules
df_sm =pd.read_csv("molecule_ID_and_SMILES.csv")
df_sm

# %% [code]
for i,row in df_sm.iterrows():
    molecule_ID = row["SAMPL6 Molecule ID"]
    smiles = row["canonical isomeric SMILES"]
    print(molecule_ID, ":", smiles)
    
    #### ENUMERATE MICROSTATES WITH EPIK ####
    # Using Schrodinger/Suites2016-4
    
    # Create the SMILES file
    !echo "Creating molecule.smi containing SMILES string..."
    #!echo "c1ccc(cc1)n2c3c(cn2)c(ncn3)N" >> molecule.smi
    with open("molecule.smi", "w") as input_file:
        input_file.write(smiles)
        input_file.write("\n")

    # Convert it to a Maestro file 
    !echo "Running ligprep to generate molecule.mae input file..."
    !$SCHRODINGER/ligprep -ismi molecule.smi -omae molecule.mae -WAIT

    # Run Epik to enumerate all "reasonable" protomers/tautomers
    !echo "Running Epik to enumerate all tautomers within 20 pK units of pH 7..."
    !$SCHRODINGER/epik -imae molecule.mae -omae epik.mae -pht 20 -ms 100 -ph 7 -WAIT

    # Extract SMILES strings for all protomers/tautomers
    !echo "Converting output to SMILES and writing epik_microstates.smi"
    !$SCHRODINGER/utilities/structconvert -imae epik.mae -osmi epik_microstates.smi

    !cp epik_microstates.smi epik_microstates.csv
    !echo "Number of predicted microstates:"
    !wc -l epik_microstates.csv

    # Create Excel file with 2D structures.
    !echo "Creating Excel spreadsheet with 2D microstate structures..."
    !python csv2xlsx.py epik_microstates.csv epik_microstates.xlsx

    !echo "Done!"
    
    
    #### Enumerate microstates with OpenEye QuacPac ####
    # Enumerate tautomers/protomers with OpenEye QuacPac
    !echo "Generating microstates with OpenEye..."

    ifs = oemolistream()
    if not ifs.open("molecule.smi"):
        OEThrow.Fatal("Unable to open SMI for reading")

    # Tautomer enumeration options
    tautomer_maxCount = 200
    tautomerOptions = OETautomerOptions(tautomer_maxCount)
    tautomerOptions.SetLevel(5)
    tautomerOptions.SetMaxSearchTime(240)
    tautomerOptions.SetCarbonHybridization(False)
    #tautomerOptions.SetRankTautomers(True)

    # Formal charge enumeration options
    charge_maxCount = 200
    chargeOptions = OEFormalChargeOptions(charge_maxCount)

    mol = OEGraphMol()

    # Unique microstate SMILES will be stored in a set
    smiles_set = set()

    while OEReadMolecule(ifs, mol):
        OERemoveFormalCharge(mol)

        # Enumerate charges first and tautomers for each charged state
        for charged_mol in OEEnumerateFormalCharges(mol, chargeOptions):
            for tautomer in OEEnumerateTautomers(charged_mol, tautomerOptions):
                smiles = OEMolToSmiles(tautomer)
                smiles_set.add(smiles) # unique SMILES are added to the set

        # Enumerate tautomers first and charges after for each tautomer
        for tautomer in OEEnumerateTautomers(mol, tautomerOptions):
            for charged_tautomer in OEEnumerateFormalCharges(tautomer, chargeOptions):
                smiles = OEMolToSmiles(charged_tautomer)
                smiles_set.add(smiles) # unique SMILES are added to the set

    with open("oe_microstates.smi", "w") as output:
        for smiles in smiles_set:
            output.write(smiles)
            output.write("\n")

    print("Done!")

    !wc -l oe_microstates.smi
    !cp oe_microstates.smi oe_microstates.csv

    # Clean up
    !trash molecule.smi

    
    #### Merge Epik and OpenEye generated microstates ####
    
    # Create a set to store Canonical Isomeric SMILES of unique microstates
    microstates_set = set()

    # Convert SMILES of Epik output to OE Canonical Isomeric SMILES

    df_epik_microstates = pd.read_csv("epik_microstates.csv", header=None)
    df_epik_microstates.columns = ["Epik output"]
    df_epik_microstates["Canonical Isomeric SMILES"] = None

    for i, row in enumerate(df_epik_microstates.iterrows()):
        smiles = row[1].values[0]

        mol = OEGraphMol()
        OESmilesToMol(mol, smiles)
        canonical_smiles = OEMolToSmiles(mol)

        df_epik_microstates.loc[i, "Canonical Isomeric SMILES"] = canonical_smiles
        microstates_set.add(canonical_smiles)


    # Convert OE output to OpenEye Canonical Isomeric SMILES

    df_oe_microstates = pd.read_csv("oe_microstates.csv", header=None)
    df_oe_microstates.columns = ["OpenEye output"]
    df_oe_microstates["Canonical Isomeric SMILES"] = None

    for i, row in enumerate(df_oe_microstates.iterrows()):
        smiles = row[1].values[0]

        mol = OEGraphMol()
        OESmilesToMol(mol, smiles)
        canonical_smiles = OEMolToSmiles(mol)

        df_oe_microstates.loc[i, "Canonical Isomeric SMILES"] = canonical_smiles
        microstates_set.add(canonical_smiles)

    # Number of microstates with unique canonical isomeric SMILES.
    # It may still include replicates due to resonance structures.
    print("Number of generated canonical isomeric SMILES (microstates + resonance str.): ", len(microstates_set))
    
    
    #### Remove duplicate resonance structures of the same microstate ####
    
    # Canonical isomeric SMILES are different for tautomers, protomers and resonance structures. 
    # We will only consider tautomers and protomers as unique microstates in this study.
    
    inchi_set = set()
    smiles_for_removal_set = set()

    for smiles in microstates_set:
        mol = OEGraphMol()
        OESmilesToMol(mol, smiles)
        inchi = prepare_inchi(mol)

        #if the same InChI is already in InChI set, drop microstate from microstates set.
        if inchi in inchi_set:
            print("Duplicate resonance structure detected. Remove:", smiles)
            smiles_for_removal_set.add(smiles)

        else:
            inchi_set.add(inchi)

    for smiles in smiles_for_removal_set:
        microstates_set.remove(smiles)

    print(len(microstates_set), " microstates were generated for ", molecule_ID, "." )
    
    
    #### Write generated microstates ####
    
    with open("{}_microstate_SMILES.smi".format(molecule_ID), "w") as output:
        for smiles in microstates_set:
            output.write(smiles)
            output.write("\n")
        
    df = pd.read_csv("{}_microstate_SMILES.smi".format(molecule_ID), header =None)
    df.columns = ["Canonical Isomeric SMILES"]
    df["Microstate ID"] = None

    for i, row in enumerate(df.iterrows()):
        id = molecule_ID+"_micro"+str(i+1).zfill(3) 
        df.loc[i, "Microstate ID"]=id

    df_microstate_ID = pd.DataFrame()
    df_microstate_ID["microstate ID"] = df["Microstate ID"] 
    df_microstate_ID["canonical isomeric SMILES"] = df["Canonical Isomeric SMILES"]
    df_microstate_ID.to_csv("{}_microstates.csv".format(molecule_ID), index=False)


    # save csv file with microstate ID and SMILES
    df.to_csv("{}_microstate_IDs_with_2D_depiction.csv".format(molecule_ID), index=False)

    # create excel file with 2D structures, microstate IDs and SMILES
    df["microstate ID"] = df["Microstate ID"]
    df["canonical isomeric SMILES"] = df["Canonical Isomeric SMILES"]

    csv_file_name="{}_microstate_IDs_with_2D_depiction.csv".format(molecule_ID)
    xlsx_file_name= "{}_microstate_IDs_with_2D_depiction.xlsx".format(molecule_ID)
    df.to_csv(csv_file_name, index=False)

    !python csv2xlsx.py $csv_file_name $xlsx_file_name
    !trash $csv_file_name
    
    # save csv for type 3 output format

    df_microstate_populations = pd.DataFrame()
    df_microstate_populations["microstate ID"]=df["microstate ID"]

    # pH range [2, 12] with 0.1 increments
    pH_range = np.arange(2,12.1,0.1)

    # make columns for each pH point
    for pH in np.nditer(pH_range):
        pH_header = "{0:.2f}".format(float(pH))
        df_microstate_populations[pH_header]=None

    df_microstate_populations.to_csv("{}_microstate_fractional_populations.csv".format(molecule_ID), index=False)

    print("Finished writing microstate files for {}!".format(molecule_ID))
    print("\n")
    print("\n")

# %% [code]
# Clean up
!trash lski*
!trash molecule.mae
!trash epik.mae
!trash epik.log
```
